# quick analysis of sedimentary ancient DNA using *quicksand*

**DOI:** 10.1101/2025.08.01.668088

**Authors:** Merlin Szymanski, Johann Visagie, Frederic Romagne, Matthias Meyer, Janet Kelso

## Abstract

Ancient DNA extracted from the sediments of archaeological sites (sedaDNA) can provide fine-grained information about the composition of past ecosystems and human site use, even in the absence of skeletal remains. However, the growing amount of available sequencing data and the nature of the data obtained from archaeological sediments pose several computational challenges; among these, the rapid and accurate taxonomic classification of sequences. While alignment-based taxonomic classifiers remain the standard in sedaDNA analysis pipelines, they are too computationally expensive for the processing of large numbers of sedaDNA sequences. In contrast, alignment-free methods offer fast classification but suffer from higher false-positive rates. To address these limits, we developed quicksand, an open-source Nextflow pipeline designed for rapid and accurate taxonomic classification of mammalian mitochondrial DNA (mtDNA) in sedaDNA samples. quicksand combines fast alignment-free classification using KrakenUniq with post-classification mapping, filtering, and ancient DNA authentication. Based on simulations and reanalyses of published datasets, we demonstrate that quicksand achieves accuracy and sensitivity comparable to or better than existing methods, while significantly reducing runtime. quicksand offers an easy workflow for large-scale screening of sedaDNA samples for archaeological research and is freely available at https://github.com/mpieva/quicksand.

## Introduction

The recovery of ancient DNA (aDNA) from archaeological sediments has enabled the retrieval of genetic material even in the absence of visible skeletal remains. Because sediments are ubiquitous at archaeological sites, extensive sample collection can be used to fill gaps in the fossil and genetic records (see Aldeias and Stahlschmidt 2024; Özdoğan et al. 2024; Evans, Llamas, and Wood 2025 for discussion). A particularly promising application is the analysis of sedimentary aDNA (sedaDNA) from the Pleistocene to gain insights into early human evolutionary history. Over the past decade, studies have shown that sedaDNA can identify the presence of ancient hominin groups (Slon et al. 2017; Braadbaart et al. 2020; Silvestrini et al. 2021), track Neanderthal and Denisovan population turnovers (Zavala et al. 2021; Vernot et al. 2021) and provide information on human subsistence strategies and human-carnivore interactions (Zavala et al. 2021; Smith et al. 2024; Gelabert et al. 2021; Gelabert et al. 2025). However, sediments present a complex mixture of DNA from microbes, plants and animals, making the recovery of ancient mammalian and human DNA particularly challenging. Hybridization capture - an enrichment method that selectively recovers DNA molecules with similarity to synthetic probes - is therefore commonly used to retrieve mitochondrial DNA (mtDNA) from a predefined set of species (Fu et al. 2013; Slon et al. 2016; Tejero et al. 2024). Because of its high cellular copy number, small genome size, and rapid rate of evolution, mtDNA is a suitable target for both enrichment and taxonomic identification. However, the accurate identification and authentication of ancient mammalian mtDNA sequences requires specialized computational workflows, as the recovered sequences are not only highly fragmented and prone to the damage-induced sequence errors typical of ancient samples (Dabney et al. 2013; Briggs et al. 2007), but also result from an unknown number of organisms and individuals.

Most workflows for sedaDNA analysis begin with the assignment of each sequence to one or more reference genomes (*classifiication*). Taxonomic classification methods can be broadly divided into alignment-based and alignment-free approaches. To date, most sedaDNA studies have relied on alignment-based classification (see SI 1) using tools such as BLAST (Altschul et al. 1990) or MEGAN/MALT (Huson et al. 2007; Herbig et al. 2016) which offer low numbers of false positive assignments (high accuracy) but require substantial computational resources (Eisenhofer and Weyrich 2019). However, the increasing size of sedaDNA datasets resulting from lower sequencing costs (Pollie 2023) and an increase in the number of available reference genomes (e.g., Sayers et al. 2023; Howard et al. 2025) have turned alignment-based methods into a major bottleneck in the analysis of sedaDNA. Alignment-free classifiers such as Kraken (Wood and Salzberg 2014) are computationally more efficient, but their use in aDNA research has been limited due to concerns over higher false positive rates (Eisenhofer and Weyrich 2019; Velsko et al. 2018; Arizmendi Cárdenas, Neuenschwander, and Malaspinas 2022). As a result, they are typically only used as a preliminary filter to reduce the dataset size before alignment (Ravishankar et al. 2024; *SediMix*: Xu, Zavala, and Moorjani 2025) or to generate a list of candidate groups for subsequent alignment-based classification (e.g. *aMeta*: Pochon et al. 2022). Studies that use alignment-free tools for taxonomic classification generally focus on the community composition rather than on verifying the individual taxonomic assignments (e.g. Margaryan et al. 2018; Courtin et al. 2022). It has been shown that taxonomic assignments generated by Kraken2 (Wood, Lu, and Langmead 2019) can be verified through additional BWA-based mapping and downstream filters (Ottoni et al. 2019; Schulte et al. 2021), highlighting the potential of combining alignment-free classification with mapping for sedaDNA analysis, but the exploration of false-positive rates and appropriate filtering strategies remains limited.

Few workflows have been described for the analysis of target-enriched mtDNA from sediments, and all rely on alignment-based classification approaches - such as the BLAST/MEGAN combination described in Slon et al. 2017 and Collin et al. 2020, competitive mapping using bwa-aln as applied in Tejero et al. 2024, or the VG Giraffe-based (Sirén et al. 2021) *vgan/euka* pipeline (Vogel et al. 2023), which aligns sequences to a pangenome graph. Of these workflows only *vgan/euka* is available as an executable tool. However, it focuses on abundance estimation and lacks several features desirable for the sensitive screening of Pleistocene sedaDNA datasets: most notably, the ability to detect low-abundance taxa (below ∼ 50 DNA sequences), to batch process input files and to customize the reference database. This highlights the need for additional user-friendly analysis pipelines that extend beyond the specific use cases supported by *vgan/euka*.

To account for these challenges, we have developed quicksand (**quick** analysis of **s**edimentary **an**cient **D**NA), a lightweight open-source Nextflow pipeline for the identification of ancient mammalian taxa from sedimentary DNA. quicksand is specifically optimized for sensitivity, speed, and portability across computing environments. It enables parallel processing of multiple sedaDNA datasets and provides options to customize the reference database. quicksand implements KrakenUniq (Breitwieser, Baker, and Salzberg 2018), a kmer-based alignment-free classifier that incorporates unique kmer-counting to improve classification specificity. To enhance its performance for aDNA we optimized KrakenUniq’s database kmer-size for the short and damaged molecules typically recovered from ancient samples (SI 3). Each KrakenUniq classification is followed by alignment with BWA (Li and Durbin 2009) and verified through post-classification and post-mapping filters to minimize false-positive rates (see SI 4). The report generated by quicksand lists the biological families identified in each sample, noting whether or not there is evidence for ancient DNA damage patterns. BAM files from each processing stage are also provided to allow further in-depth analysis with external tools.

## Results and Discussion

### Workflow

#### Reference Genomes and Database

Before quicksand is run, reference-files and the KrakenUniq kmer-database must be constructed from the set of reference genomes to be included in the analysis. This step is handled by the quicksand helper pipeline ‘quicksand-build’ (see ‘Availability’ section). By default ‘quicksand-build’ includes all mtDNA reference genomes in the NCBI RefSeq database (O’Leary et al. 2015). NCBI RefSeq provides high-quality, non-redundant reference genomes for more than 1800 mammalian species that are linked with the NCBI taxonomy used by KrakenUniq for classification. All genomes are indexed with BWA and low complexity regions in the mtDNA sequences are masked using ‘dustmasker’ (Morgulis et al. 2006).

#### Data Preprocessing

The input for quicksand is a directory with user-supplied DNA sequence files in BAM or FASTQ format. Demultiplexing, overlap merging and adapter-trimming need to be performed by the user prior to running quicksand.

#### Running quicksand

quicksand analysis proceeds through four steps (Fig. 1):

1. Taxonomic classification and binning of individual DNA sequences with KrakenUniq
2. Mapping of binned sequences with BWA and removal of PCR duplicates.
3. Analysis of deamination signals on a per-family basis.
4. Calculation of summary-statistics for filtering false-positive assignments

**Figure 1.**
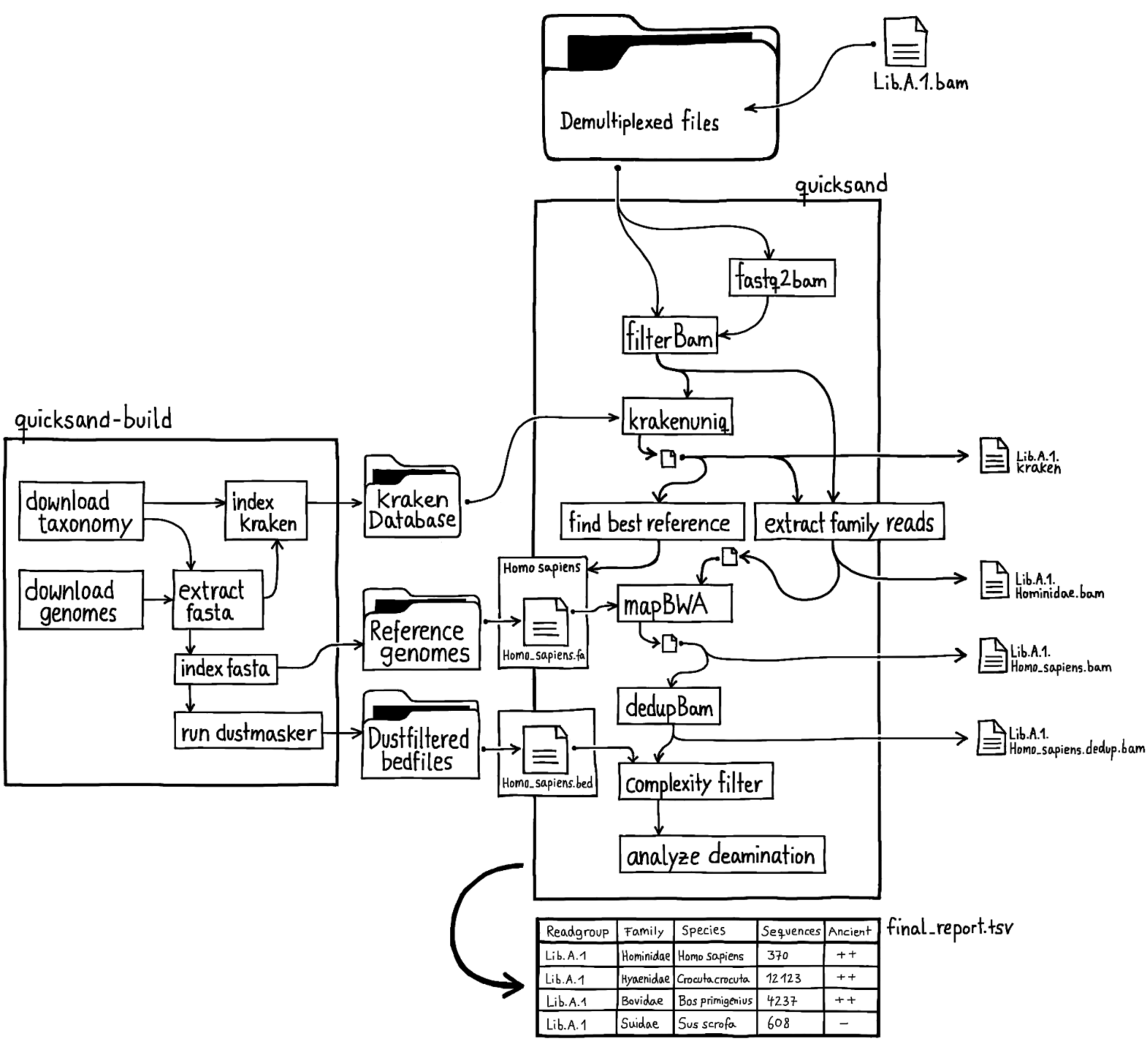
Schematic overview of the quicksand workflow. The file names are examples for classification of sequences as Hominidae. The“quicksand-build” panel shows the workflow of the quicksand-build pipeline for creating the required reference data-files.

### 1. Taxonomic Classification and Binning

Only sequences that are long enough for reliable classification and mapping (default: >= 35 basepairs) are classified by KrakenUniq. Sequences are then binned by biological family or biological order (default: family) and bins with too few sequences (default: < 3) or too few unique kmers (default: < 129, see SI 4) are removed from further analysis. For each identified family, the reference genomes from the node with the highest number of unique kmers are selected for mapping. To find this node, quicksand implements a naive decision tree, following the path of the highest kmer counts in the taxonomic tree from the family to the species level (see Table 1).

**Table 1.** Example for the selection of a reference genomes.

| Rank | Taxa | Kmers | Selected |
| --- | --- | --- | --- |
| family | Hominidae | 400 |  |
| genus | \|-- Homo | 300 | X |
| genus | +-- Pan | 5 |  |
| species | +-- Pan troglodytes | 2 |  |

The table shows a simplified KrakenUniq report. The "Selected" column indicates the node selected by quicksand for the mapping step. In this example, the Hominidae (great apes) family is classified by 400 unique kmers. Of these, 300 match the *Homo* (humans) genus and five match the *Pan* (chimpanzees) genus. Although two kmers match the *Pan troglodytes* species level, the majority of the genus-level kmers fall under *Homo*, which is therefore selected as the reference node for the Hominidae family

In the example above (Table 1), the genus *Homo* is selected as reference-node for the Hominidae family. The mtDNA reference genomes used for mapping are therefore those contained within this family in NCBI RefSeq (*Homo sapiens* (NC_012920.1), *Homo neanderthalensis* (NC_011137.1), *Homo sapiens subsp. Denisova* (NC_013993.1) and *Homo heidelbergensis* (NC_023100.1)).

### 2. Mapping and De-duplication

The sequences assigned to each family are then mapped to all selected reference genomes. Mapping is performed using the “network awareˮ fork of BWA, version 0.5.10-evan.10 (https://github.com/mpieva/network-aware-bwa/) with ancient parameters (-n 0.01-o 2-l 16500) (Meyer et al. 2012). Unmapped sequences and those below the mapping quality threshold (default: 25) are removed from the alignment using samtools v1.15 (Danecek et al. 2021). If multiple reference genomes were used (as in the *Homo* example above), a single “bestˮ reference genome is kept based on the highest number of covered base pairs in the alignment. Sequences with identical alignment start and end coordinates are then collapsed into a unique DNA consensus sequence using bam-rmdup (https://github.com/mpieva/biohazard-tools) v0.2.2. In a final step, to reduce spurious alignments, sequences that overlap low-complexity regions in the reference genome by at least one base pair are discarded using bedtools intersect (Quinlan and Hall 2010).

### 3. Determining Signals of Deamination

Cytosine deamination, which results in the conversion of cytosines (C) to uracils (U), particularly at the terminal positions of DNA strands, is the most common miscoding lesion seen in ancient DNA. When sequenced, uracil is misread as thymine (T), creating C-to-T differences to the reference genome, which are characteristic of aDNA sequences. To assess whether ancient DNA is present, quicksand computes C-to-T substitution frequencies at the 5’ and 3’ termini of sequences assigned to each biological family, together with their 95% binomial confidence intervals. Following the strategy described in Slon et al. 2017, these values are then used to report families containing authentic aDNA sequences in the ‘Ancientness’ column of the summary report file. The significance of the deamination signals are indicated as follows: (++) C-to-T substitutions frequencies on both ends of sequences are significantly higher than 10% (i.e., the lower limit of the 95% binomial confidence interval exceeds 10%). Sequences in this category are considered to originate at least partly from authentic aDNA.

(+) C-to-T substitution frequencies on either the 5’ or the 3’ end of sequences are significantly higher than 10%, requiring further evaluation. These values could result from low to medium damage, as expected for instance from Holocene samples, or a lack of power in detecting authentic older aDNA.

(-) C-to-T substitutions frequencies at neither the 5’ nor the 3’ end significantly exceed 10%, indicating that there is no sufficient evidence for the presence of aDNA from the respective family.

### 4. Post-pipeline Filtering

False positive identifications are a well known problem for metagenomic analyses and can lead to incorrect conclusions (e.g., see Haas et al. 2022 in reply to Weyrich et al. 2017). Workflow-specific filters should therefore be implemented to minimize false positive classifications. To facilitate such filtering, quicksand provides summary statistics that allow for additional filtering of detected families based on relative sequence quantity (percentage of unique sequences per family, “PSFˮ) as well as the evenness of sequence coverage of the reference genome (proportion of expected breadth, “PEBˮ; i.e. the observed breadth of coverage / expected breadth of coverage). Based on our testing (see SI 4), we recommend removing families that are supported by a PSF of less than 0.5%, as well as families with a PEB below 0.5. In addition to the full report, quicksand provides a pre-filtered version of the table with these thresholds already applied.

### Customization

quicksand provides several options (*filags*) for customization, which are documented in detail on readthedocs (https://quicksand.readthedocs.io/en/latest/). While most available flags adjust the filter thresholds described above, two flags have an impact on the data processing workflow and are therefore briefly outlined below. The first option is the choice of the classification level used for taxonomic binning, which is set at either the biological family level (default) or order level. The second option allows specific, pre-defined reference genomes (“fixed references”) to be selected for mapping after KrakenUniq classification.

### Binning by Order-level

By default quicksand uses the KrakenUniq classification to bin sequences by biological family before mapping them to the corresponding “best” family-level reference genome(s) as described above. To increase sensitivity, especially for sequences that are strongly divergent from the available reference genomes and may thus be misassigned at the family level, it is possible to set the binning to the biological order level. In this case, all sequences assigned to an order (e.g., Primates) are binned before being mapped to the specific family-level reference genomes selected earlier (e.g., *Homo* for Hominidae sequences).

### Mapping to Pre-selected Reference Genomes

In a default run, sequences are mapped to only a single reference genome per detected family. However, quicksand allows users to define one or more specific reference genomes per family instead. This option was implemented to enable downstream analyses that depend on specific alignments - such as the human mtDNA haplotype caller mixEMT (Vohr et al. 2017), which requires mapping of human sequences to the Cambridge Reference Sequence (rCRS; NCBI Accession: NC_012920.1; Andrews et al. 1999). When such predefined (“fixedˮ) references are used, quicksand adds an additional workflow step, saving the putatively deaminated sequences in a separate BAM file for potential later use.

### Optimization of KrakenUniq Kmer-size for aDNA

KrakenUniq assigns each DNA sequence to a node in a pre-defined taxonomic tree by splitting the DNA sequence into overlapping fragments of length k (kmers) that are then each perfectly matched to a pre-indexed kmer database. The default kmer size for KrakenUniq is 31. However, for aDNA, classification by perfectly matching kmers is complicated by the short length of the sequences, terminal base substitutions due to deamination, and sequence divergence between the ancient species in the sample and the modern reference genomes used for classification. For these reasons, we hypothesized that a shorter kmer size would be better suited for the classification of aDNA, as more kmers from the middle part of the sequences, which are less affected by deamination, can be taken into account, making it more likely to match a kmer-reference in the kmer database.

To verify this hypothesis, we constructed kmer-databases indexed with sizes 18, 19, 20, 21, 22, 24, 28 and 30. These databases included the 1556 mammalian mtDNA reference genomes in NCBI RefSeq Release 218. We then used gargammel (Renaud et al. 2016) to simulate 15 paired-end Illumina MiSeq datasets. These datasets contained 500 to 5,000,000 mammalian mtDNA sequences (dataset size), as well as mammalian nuclear DNA and environmental “backgroundˮ DNA (DNA from bacteria, archaea, fungi, plants, and viruses), totaling between 500,000 and 5,000,000 sequences each (see SI 2). To mimic authentic aDNA, the length of the simulated sequences followed the read length distribution of a shotgun-sequenced aDNA library from the *Mezmaiskaya 2* Neanderthal bone (A9180, Prüfer et al. 2017). In addition, varying levels of deamination damage were introduced at the ends of the reads, following two published deamination profiles (medium: Olalde et al. 2014, high: Günther et al. 2015). Nine of the 15 datasets included the environmental background DNA to evaluate its effect on sequence classification. Sequences were post-processed to remove sequencing adapters and overlapping read pairs were merged into full-length aDNA molecules using leeHom (Renaud, Stenzel, and Kelso 2014). Finally, all datasets were analysed with quicksand v2.3 using each of the constructed kmer-databases to find the kmer-size best suited for sedaDNA datasets.

### Assignment Accuracy

We calculated the assignment accuracy as the proportion of correctly assigned sequences (true positives) to the total number of assigned sequences (true positives + false positives). We found that assignment accuracy negatively correlates with kmer size, magnitude of DNA damage, and dataset size (Fig.2). In datasets without simulated damage, the median accuracy ranged from 98.96% (kmer 18) to 99.98% (kmer 21–30), with minimum values between 93.18% and 99.42%. In contrast, in medium and high damage datasets median accuracy ranged from 98.66% and 96.08% (kmer 18) to 99.76% and 98.69% (kmer 30), respectively. Outliers were observed in the largest datasets with 5 million input sequences (∼94% for medium damage and ∼86% for high damage; Fig.2A). We observed that lower accuracies were primarily driven by the increased number of false-positive family assignments (see SI 3 and 4) but that applying the recommended post-pipeline filters on the family-level (0.5% PSF and 0.5 PEB, see SI4) increased median accuracy across all datasets and kmer sizes to above 99.75% (Fig.2B). This demonstrates that even at lower kmer sizes, post-processing can limit the number of false positive assignments.

**Figure 2.**
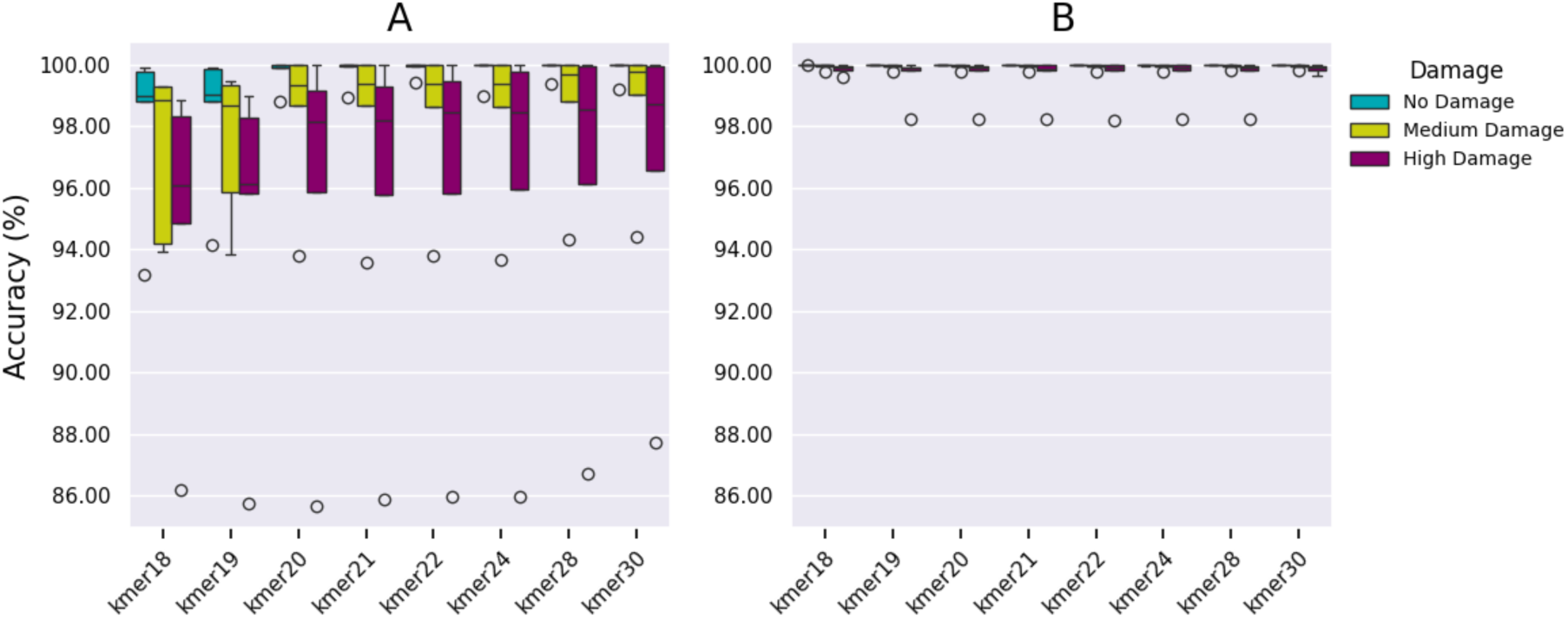
Assignment accuracy per kmer-size in (A) unfiltered quicksand results and (B) with applied post-pipeline-filters on the family level (≥0.5% of sequences and PEB ≥0.5, see SI 4), stratified by the damage-profile of the simulated datasets. The boxplot shows the accuracy distribution across 15 datasets (5 per box). Box limits represent the 25th and 75th percentiles, the center line indicates the median, and whiskers extend 1.5 times the interquartile range. Outliers are shown as dots.

### Assignment Sensitivity

To assess which kmer size yields the highest sensitivity (i.e., the highest number of correctly identified sequences), we calculated the number of correctly assigned sequences per dataset and family relative to the *highest* number obtained for that dataset and family across all kmer sizes (Relative Sensitivity). Fig.3 shows the distribution of the relative sensitivity scores for (A) no damage, (B) medium damage and (C) high damage datasets. In the no-damage and medium-damage datasets, the number of sequences recovered correlates with increasing kmer-sizes, peaking at kmer-size 30. In contrast, in the highly damage datasets, sensitivity is highest for kmer-sizes 19 to 24.

**Figure 3.**
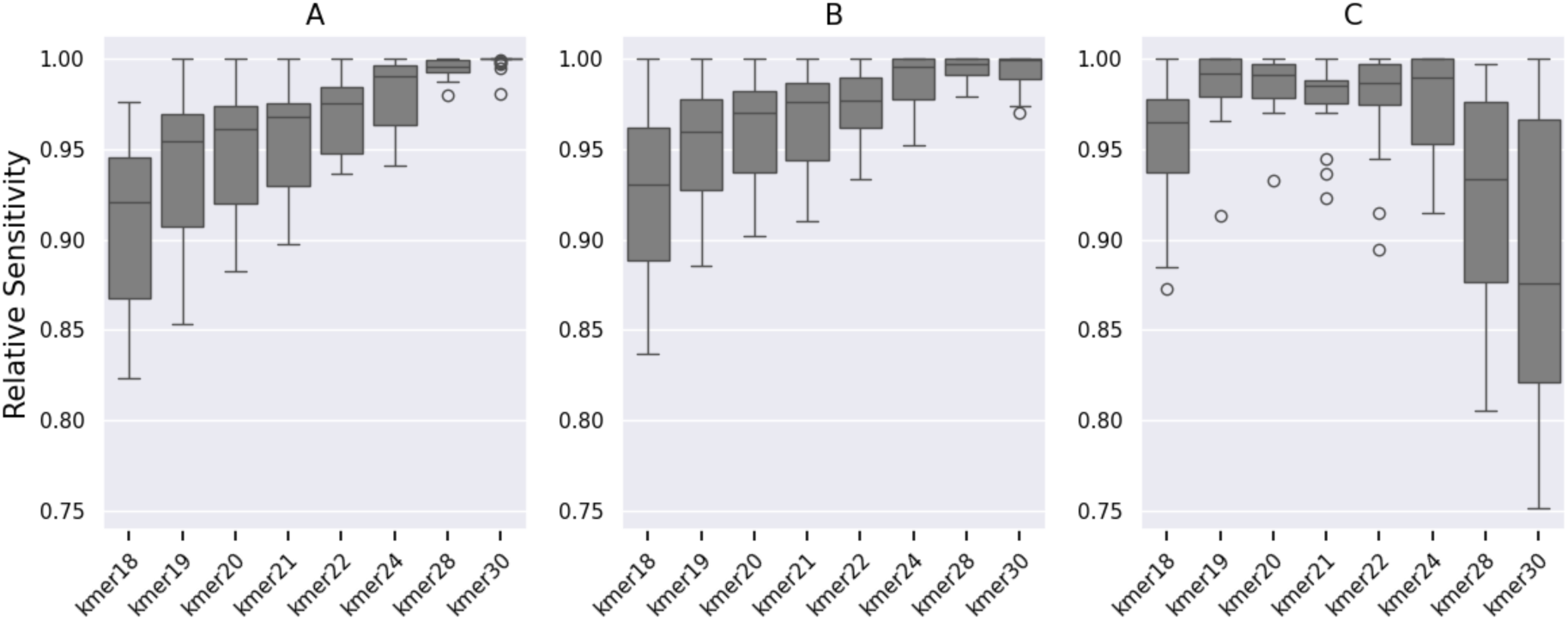
Relative sensitivity per kmer size in (A) no damage, (B) medium damage, (C) high damage datasets. Each boxplot shows the distribution of relative sensitivities per datasets and biological family within each damage category (25 data points per box). Box limits represent the 25th and 75th percentiles, the center line indicates the median, and whiskers extend 1.5 times the interquartile range. Outliers are shown as dots.

With the accuracy remaining stable across all kmer-sizes after applying post-pipeline filters (Fig. 2) and sensitivity increasing for kmer sizes 19-24 in the highly damaged datasets, we decided on a kmer-size of 22 as the default for quicksand. For samples with less damage, the kmer-size could be adjusted to 24 or higher (e.g. for Holocene samples) while for more damaged samples the kmer size could be reduced to 19 or 20.

### Benchmarking

We used the simulated datasets described above to benchmark quicksand v2.3 against both the BLAST/MEGAN-based workflow described in Slon et al. 2017 and *vgan/euka* (Vogel et al. 2023), comparing assignment accuracy, sensitivity and runtime. To simplify comparison we implemented the BLAST/MEGAN and *vgan/euka* methods as Nextflow pipelines (see SI 5), enabling them to process the same input format and produce summary reports comparable to that of quicksand. All three pipelines were executed on the same compute node with 64 CPUs and 1056.63 GB memory.

### 1. Accuracy

We counted positive sequence assignments on the level of biological families. However, for *vgan/euka,* we considered sequences to be true positives if they were assigned to the correct pangenome sub-graphs (i.e., Carnivora for hyaena, Proboscidea for mammoth, Suina for pigs, Bovidae for bovids, Homininae for humans; see SI 5). As shown in Fig.4A, both quicksand and BLAST/MEGAN consistently achieved median accuracies above 99.5%. In contrast, *vgan/euka’s* accuracy declined as dataset size increased, dropping from a median of above 99% at 500 mtDNA sequences to approximately 95% in the largest dataset containing 5 million mtDNA sequences. As shown in Table 2 this is explained by a larger number of false positive assignments with increasing sequence input.

**Figure 4.**
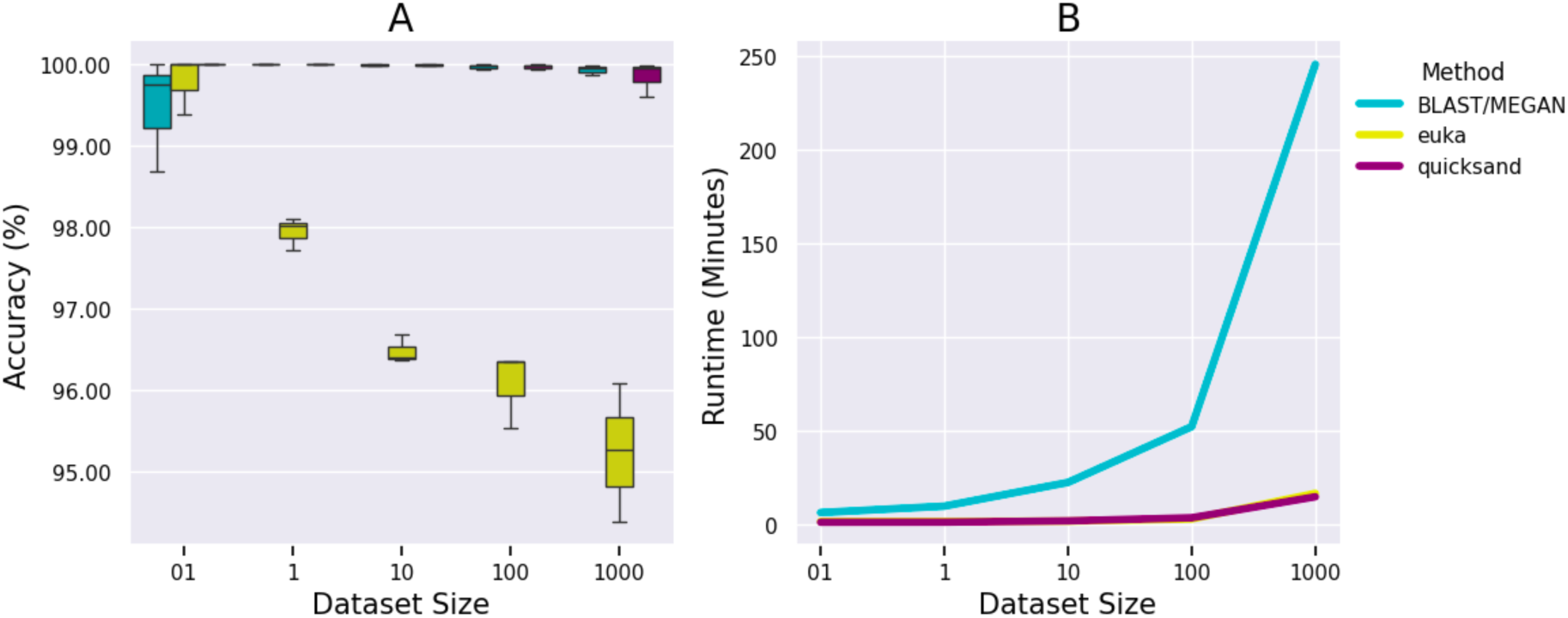
Benchmarking *quicksand* against the BLAST/MEGAN and *vgan/euka* pipelines. Dataset sizes are described in SI 2 and range from 500 to 5 million mtDNA sequences per dataset. (A) Assignment accuracy. Each box represents the accuracies obtained for the datasets within a given size category (3 datasets per box). Box limits represent the 25th and 75th percentiles, the center line indicates the median, and whiskers extend 1.5 times the interquartile range. Outliers are shown as dots. (B) Pipeline runtime in minutes.

**Table 2.**
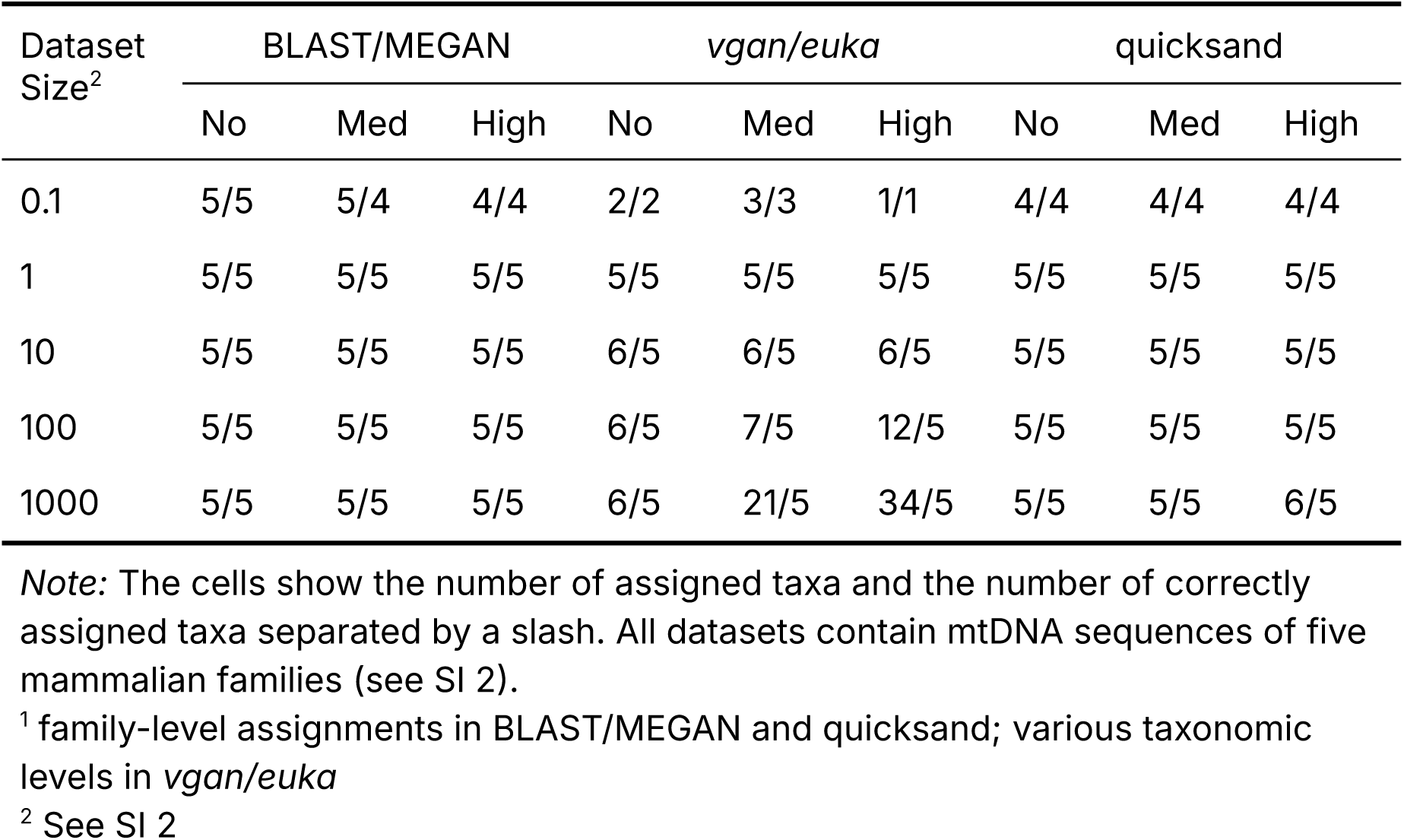
Assigned taxa^1^ per simulated dataset, damage profile and method.

### 2. Sensitivity

While quicksand and BLAST/MEGAN include a ‘bam-rmdup’ step to remove PCR duplicates after mapping, *vgan/euka* does not perform sequence deduplication, making it impossible to compare counts of correctly identified sequences among these methods. Table 2 therefore provides an overview of the number of taxa identified with the three methods. We find that both BLAST/MEGAN and quicksand are more sensitive than *vgan/euka* in the smallest datasets, consistently detecting four out of the five families, based on as few as 20 mtDNA sequences. In contrast, *vgan/euka* only found one to three families in the smallest datasets (the ones with the highest number of sequences), consistent with the reported requirement of ∼50 sequences for a successful detection (Vogel et al. 2023). BLAST/MEGAN was the only method to successfully detect the presence of humans in the smallest dataset based on just five (undamaged) mtDNA sequences (see SI 4.1 and SI 6).

### 3. Runtime

We grouped the 15 simulated datasets into five categories based on the dataset size (3 BAM files per group) and measured the runtime of each group and pipeline using the unix time command. Both quicksand and *vgan/euka* were similarly fast, with runtimes between 1 minute for the smallest datasets and 14 and 16 minutes, respectively, for the largest. In contrast, the BLAST/MEGAN pipeline showed a much steeper increase in runtime, taking 6 minutes for the smallest dataset and 245 minutes for the largest (Fig.4B).

Overall, quicksand performs well across all three benchmarking criteria. It matches the BLAST/MEGAN pipeline in terms of accuracy and sensitivity, while outperforming it in runtime. Compared to *vgan/euka*, quicksand maintains high accuracy even with large dataset sizes, requires fewer mtDNA sequences for successful family-level detection and provides finer taxonomic resolution - all while achieving comparable processing speeds.

### Verification of quicksand using real data

To evaluate the performance of quicksand v2.3 on real data, we re-processed a published dataset, consisting of 274 Pleistocene sediment DNA libraries from the Main Chamber of Denisova Cave, Russia, enriched for both mammalian mtDNA (probe set AA75) as well as specifically hominin mtDNA (probe set AA163; see Zavala et al. 2021, Supplementary Data 1, “Mammalian MtDNA Summaryˮ and “Hominin MtDNA all Libˮ; “First Screenˮ libraries). For each of the samples, between 1.5 and 3 million reads were generated on the Illumina HiSeq2500 platform and analyzed using the BLAST/MEGAN-based approach. Overall, the dataset combines multiple challenging features, such as large numbers of sequences generated per sample, high C-to-T deamination frequencies (up to 75% at the ends of molecules) and ‘background’ DNA of unknown composition.

Libraries captured with the mammalian mtDNA probes were analyzed using quicksand’s default parameters. We found that applying the recommended post-pipeline filters (PSF of at least 0.5% and PEB of at least 0.5) effectively removed the majority of putative false-positive family assignments (see SI 7.1). Consistent with the workflow described by Zavala et al. (2021), libraries captured with human mtDNA probes were analyzed using the “fixedˮ option, ensuring that sequences classified as Hominidae were aligned to the rCRS for additional downstream analyses. No PSF filter was applied to those libraries.

quicksand identified ancient mammalian mtDNA in 271 samples, compared to 267 samples reported by Zavala et al. 2021, and recovered a larger number of unique sequences per biological family (Fig. 7A). Sequences from additional ancient families were detected in 205 samples (Table SI7.1), most often from Hyaenidae, which were found in 58 additional samples and are dominant in nearly all samples from the upper layers (layer 19 and above; Fig. 5) in the quicksand analysis. Across all samples, a total of 13 ancient families were identified. These families are identical to those previously reported by Zavala et al. 2021. Apart from the increased abundance of hyaena mtDNA, the composition of ancient mammalian families per sample closely resembles earlier findings (Fig. 5), with bear and canid mtDNA dominating the lower layers (22.1 and 20) and a shift toward bovid and cervid mtDNA in the upper layers.

**Figure 5.**
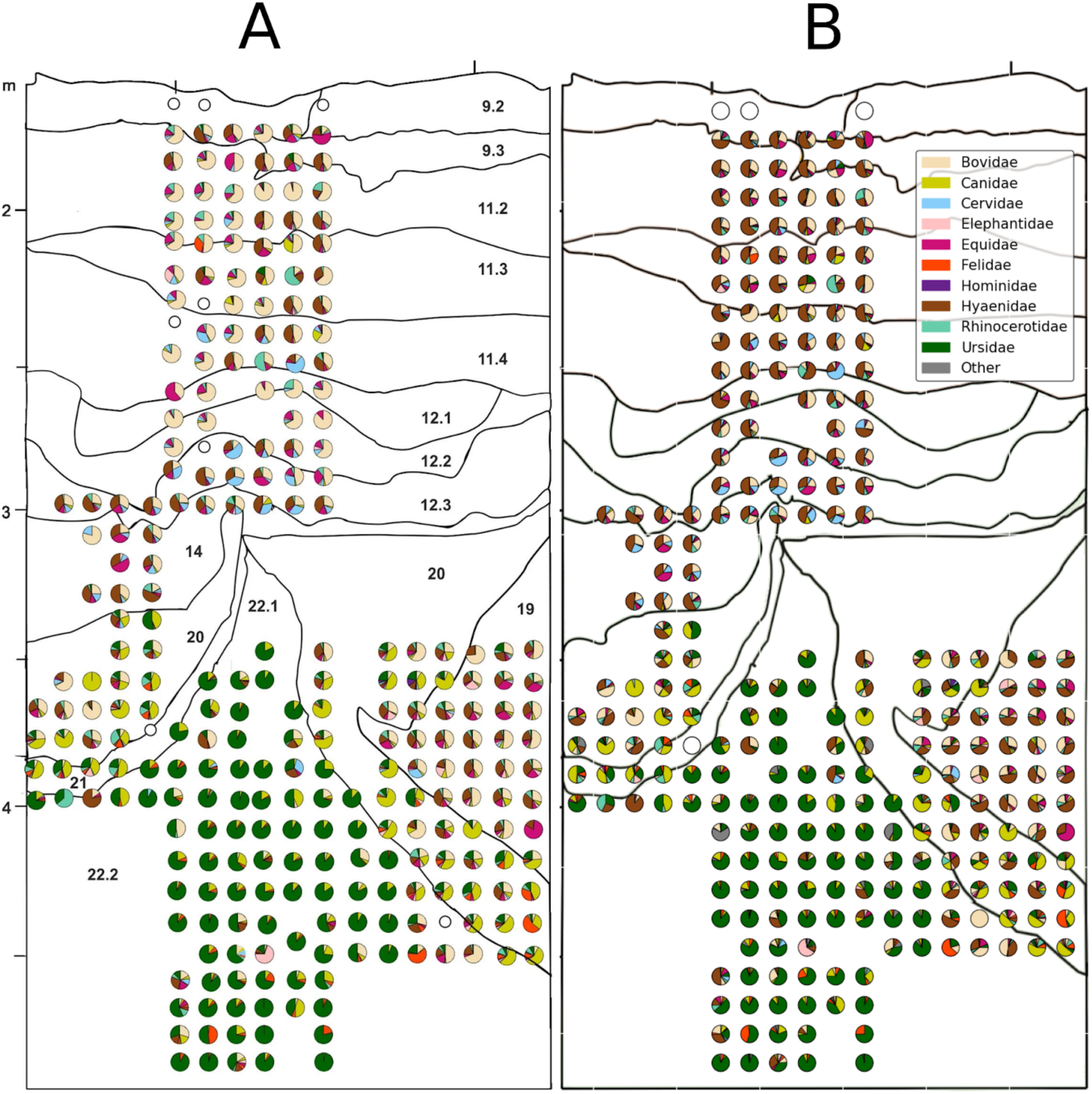
Composition of ancient mammalian mtDNA in the Main Chamber of Denisova Cave. The pie charts display the proportions of unique mtDNA fragments assigned to each ancient mammalian family in the sample. White circles indicate samples with no evidence for ancient mtDNA. (A) Extended Data Fig.8, as published in Zavala et al. 2021 (B) Recreation of panel A using the results from the quicksand analysis.

**Figure 6.**
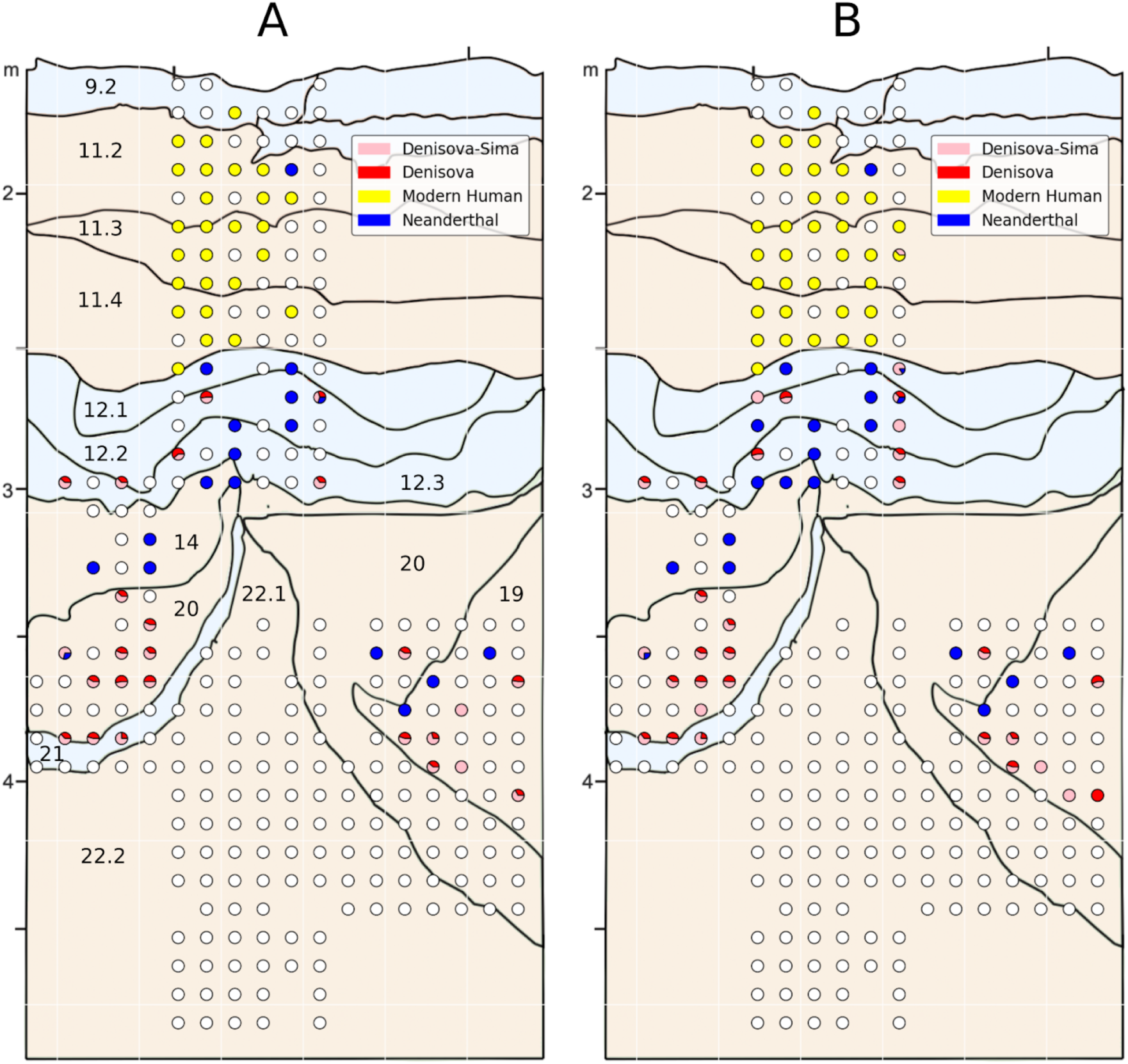
Composition of hominin mtDNA lineages detected in samples from Denisova Cave, Main Chamber. The pie charts display the proportion of mtDNA fragments assigned to the different hominin lineages. White circles indicate samples with no evidence for ancient human mtDNA or with less than 10% support for any given lineage. (A) Plot created from data published in Zavala et al. 2021 (Supplementary Data 1, “Hominin MtDNA all Libˮ all “First Screenˮ libraries) for comparison (B) the same figure, created using the results from quicksand.

**Figure 7.**
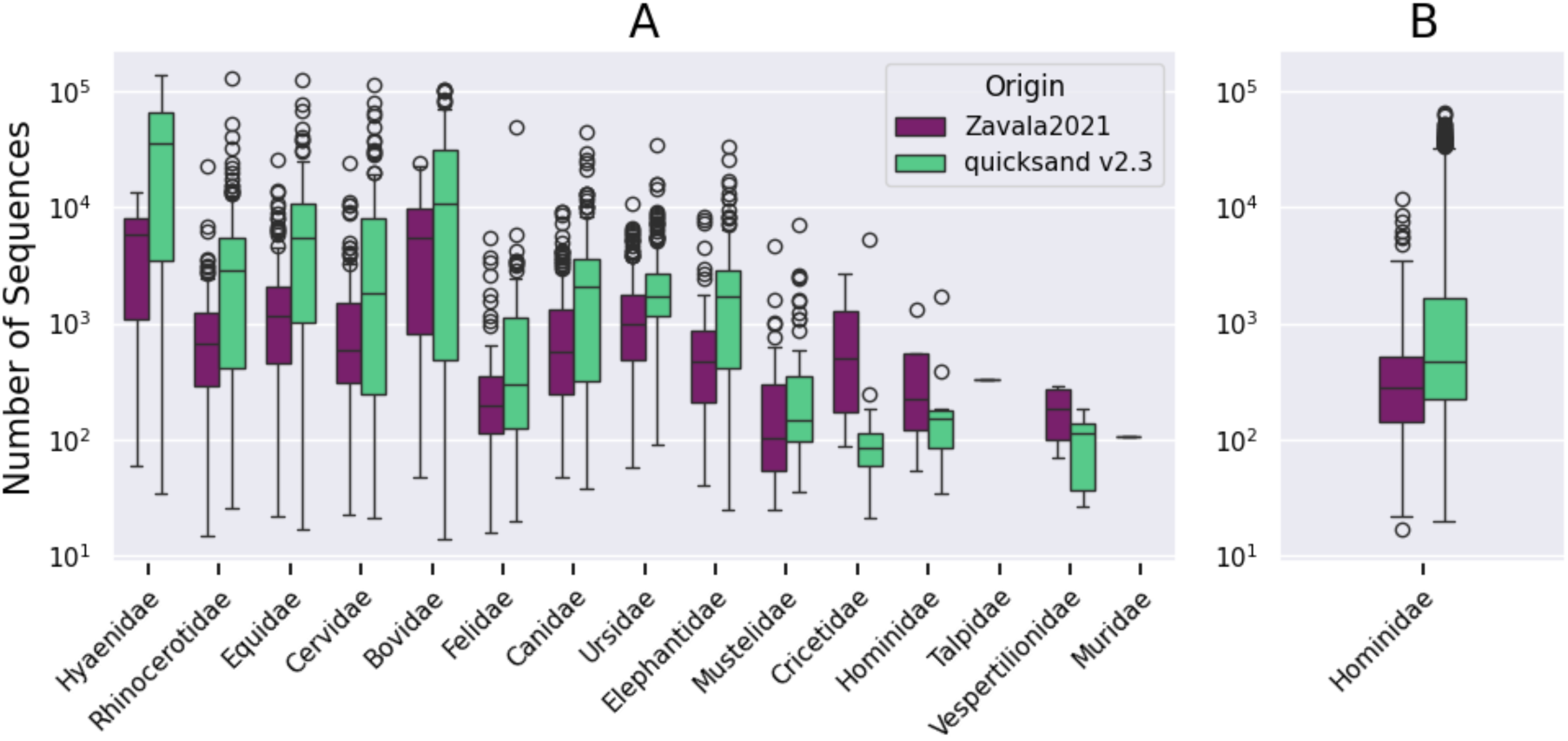
Number of unique mtDNA sequences recovered for each ancient mammalian family. Data as published in Zavala et al. 2021 or analyzed with quicksand v2.3 (A) across the 274 mammalian capture libraries (B) across the 274 human capture libraries. Box limits represent the 25th and 75th percentiles, the center line indicates the median, and whiskers extend 1.5 times the interquartile range. Outliers are shown as dots.

quicksand also showed a higher success rate in the identification of ancient hominin DNA (Fig. 7B), with 100 samples yielding a positive result compared to 77 with BLAST/MEGAN. Only one sample dropped from firm evidence (++) to suggestive evidence of ancient hominin DNA (+), due to a slight decrease in the sample’s terminal deamination frequency. In a lineage assignment test, which calculates the support for different branches in the hominin mtDNA tree (Neanderthals, Sima de los Huesos hominins, Denisovans, and modern humans; see SI 7.2 for details), the distribution of the identified lineages in the stratigraphy (Fig.6) corresponds to the one described by Zavala et al. 2021. No hominin mtDNA was recovered from layer 22 and the oldest mtDNA fragments were assigned to the Denisovan lineage. As reported before, we find a mixture of Denisovan and Neanderthal mtDNA in samples from the middle Middle Palaeolithic (mMP) layers 12-19 and a lack of Denisovan mtDNA in layer 14, consistent with an occupation by Neanderthals only during the formation of this layer. Modern humans are exclusively present in samples associated with the Initial Upper Palaeolithic (IUP) industries from layer 11 to layer 9.

In summary, for both the mammalian and human mtDNA captured libraries, we successfully replicated the results of the BLAST/MEGAN based workflow using quicksand and demonstrate that quicksand recovers more sequences on average from all ancient families, providing more power for downstream analyses. We also demonstrate that the filtering criteria developed using simulated data are suitable for the application of quicksand to real data.

### Pipeline Portability and Data Availability

quicksand is developed in the Nextflow programming language (Di Tommaso et al. 2017) following nf-core community standards (Ewels et al. 2020) of code structure and modularization. For independence of the computational platform and installed software versions, quicksand runs its processes in software-containers (*images*) using Docker (Merkel 2014) or Singularity (Kurtzer, Sochat, and Bauer 2017). Software images are pulled on demand from Dockerhub (https://hub.docker.com/u/merszym), the Galaxy Project singularity image depot (https://depot.galaxyproject.org/singularity/) or the quai.io biocontainers repository (https://quay.io/organization/biocontainers). For custom software, Dockerfiles and images were created by the authors and stored in the pipeline git repository and on Dockerhub respectively.

**quicksand** is available under MIT License on *github* (https://github.com/mpieva/quicksand) with detailed documentation on *readthedocs* (https://quicksand.readthedocs.io/).

The most recent pre-constructed version of the quicksand **database** can be downloaded from the MPI EVA *FTP server* (http://ftp.eva.mpg.de/quicksand/build/).

quicksand-build offers multiple options for customization, which includes adding custom mtDNA genomes to the database or using a customized version of the NCBI taxonomy.

**quicksand-build** is available under MIT License on *github*

(https://github.com/mpieva/quicksand-build).

The python-notebooks used to create the figures in this paper can be found on *github*

(https://github.com/merszym/quicksand-paper-analyses)

### Hardware requirements

Using the provided pre-computed reference-databases described above (57GB), quicksand can be run on a standard personal laptop (e.g. 11th Gen Intel(R) Core(TM) i7-1165G7 @ 2.80GHz with 8 threads, 32GB RAM) due to the memory-efficient database loading implemented in KrakenUniq >= v1.0.0 (Pockrandt, Zimin, and Salzberg 2022) and the relatively small size of the NCBI RefSeq mtDNA database (∼50GB).

For the construction of a custom database with ‘quicksand-build’, a dedicated machine with at least 100GB of RAM is required.

## Conclusion

We present quicksand, an efficient and portable bioinformatics pipeline specifically optimized and validated for the identification and taxonomic classification of ancient mammalian mitochondrial DNA (mtDNA) from sediment samples. quicksand combines alignment-free classification with downstream BWA alignments and post-classification filtering to increase classification speed while maintaining high accuracy. Using simulated data, we demonstrate that a kmer size of 22 accounts for the short fragment lengths and deamination damage patterns characteristic of aDNA datasets. We validate quicksand with simulated datasets and with sedaDNA data from Denisova Cave, successfully replicating previously published results and detecting additional samples positive for ancient mammalian and hominin mtDNA.

Benchmarking analyses show that quicksand matches alignment-based approaches such as BLAST/MEGAN in terms of accuracy and sensitivity, while offering substantial improvements in runtime and portability. Compared to *vgan/euka*, quicksand maintains higher assignment accuracy across the full range of dataset sizes and performs better with small numbers of sequences. Overall, quicksand offers a fast and easy-to-use alternative to alignment-based pipelines for sedaDNA research, enabling efficient screening of large datasets. While quicksand was developed and optimized for the detection of ancient mammalian mtDNA in sediments, its database customization options make it adaptable to a wider range of use cases, such as the detection of additional taxa in sediment DNA data or the analysis of other metagenomic datasets.

## Acknowledgements

This project was funded by the Max Planck Society

