## Supplementary Information for "quick analysis of sedimentary ancient DNA using *quicksand*"

#### Supplementary Section 1: Published Workflows in Ancient Metagenomics and sedaDNA Research

The term *ancient metagenomics* refers to the analysis of shotgun-sequenced samples for ancient DNA (see Fellows Yates et al. 2024) to emphasize the focus on multiple (unknown) sources of the DNA in the sample and the untargeted nature of shotgun-sequencing. In this section we summarize bioinformatic workflows commonly used in the field of sedaDNA or ancient metagenomics that were considered during the development of quicksand.

In the main text we focused on the advances of sedaDNA in the field of archaeology. However, the first recovery of sedaDNA came from sediment-cores used for the *reconstruction of past ecosystems* (e.g. Willerslev et al. 2003; Hofreiter et al. 2003). Since then, shotgun sequencing of sedaDNA has been widely applied to permafrost, marine, and lake sediments to investigate past biodiversity and palaeoenvironments (e.g. Kjær et al. 2022; Wang et al. 2021; Capo et al. 2021; Parducci, Nota, and Wood 2018; Pedersen et al. 2016; Liu et al. 2024). Bioinformatic workflows for ancient metagenomic data are generally agnostic to the sample type, and have been used across a wide range of substrates. In addition to workflows developed specifically for sedaDNA analysis, we also explored methods focused on ancient microbial communities - for example, those targeting host-associated microbiomes or pathogens in archaeological materials. The computational workflows generally involve:

(i) taxonomic classification, where individual DNA sequences are assigned to one or multiple reference genomes and placed on a phylogenetic tree. (ii) taxonomic binning, where sequences from the same taxonomic level (e.g., biological family) are grouped for downstream analyses. (iii) data filtering and validation. (iv) Analysis of DNA damage patterns to confirm sequence authenticity.

For the development of quicksand, we compared bioinformatic workflows and their classifiers and evaluated strategies for the authentication of results and the removal of false positives. In particular, we screened workflows published in the following fields of ancient DNA research:

1. Sediments, environmental reconstruction
2. Sediments, archaeology
3. Other substrates, (host-associated) microbial communities

In addition, we specifically surveyed aDNA studies that used tools from the Kraken family (Kraken, Kraken2 or KrakenUniq) for taxonomic classification to better understand how false-positive assignments can be addressed and removed during downstream analyses.

**Table S1.1**
Overview of workflows in sedaDNA and ancient metagenomics studies

| Name / Publication | Classifier | Main Filters | Substrate / Method |
|---|---|---|---|
| HOLI (Pedersen et al. 2016) | Bowtie2 + MEGAN or ngsLCA | Minimum number of sequences; Minimum percentage of sequences | SG from sediments; environmental Reconstruction |
| Slon et al. 2017 | BLAST + MEGAN. Additional mapping with BWA | Minimum number of sequences; Minimum percentage of sequences | TE from sediments, (Target: mammalian and hominin mtDNA); Archaeology |
| Margaryan et al. 2018 | **Kraken** | Minimum Kraken confidence score; Minimum number of sequences; | SG from other substrate; Host-Associated Microbial Communities |
| HOPS (Hübler et al. 2019) | MALT | Minimum sequence complexity; Shape of edit distance; Ancient damage patterns | SG and TE, all substrates; Microbial Communities |
| Ottoni et al. 2019 | **Kraken2**. Additional mapping with BWA | Minimum percentage of sequences; Species blacklist; | SG from other substrate; Host-Associated Microbial Communities |
| "An open sourced pipeline" (Collin et al. 2020) | BLAST + MEGAN. Additional mapping with BWA | Minimum number of sequences; Minimum percentage of sequences | SG from sediments; Archaeology |
| Schulte et al. 2021 | **Kraken2**. Additional mapping with BWA | Kraken confidence score; | TE from sediments (Target: *Larix* Chloroplast); environmental Reconstruction |
| Courtin et al. 2022 | **Kraken2** | Kraken confidence score; Minimum percentage of | SG from sediments; environmental |

| | | | |
|---|---|---|---|
| | | sequences;<br>Assignment to “likely” family | Reconstruction |
| HAYSTAC (Dimopoulos et al. 2022) | Bowtie2 | Minimum breadth of coverage;<br>Minimum posterior probability | SG from all substrates; Microbial Communities |
| aMeta (Pochon et al. 2022) | Profiling with **KrakenUniq**, then MALT | Minimum number of kmers;<br>Minimum number of sequences | SG from all substrates; Microbial Communities |
| *vgan/euka* (Vogel et al. 2023) | VG Giraffe | Spurious alignment filter;<br>Minimum coverage | SG/TE from sediments (Target: tetrapod and arthropod mtDNA); Archaeology |
| sediMix (Xu, Zavala, and Moorjani 2025) | **CENTRIFUGE**. Additional mapping with BWA | Required assignment to Primates; | TE from sediments, (Target: Human nuclear DNA); Archaeology |

*Notes*: The table focuses on the classifier and filter steps of published workflows. Workflows *without* a name are only described in the respective publication and not available as an executable tool or pipeline. The “Main Filters” column shows a selection of filters applied on the classified sequences to remove false-positive assignments. See the respective publications for more details.

Alignment-free classifiers in bold
*Abbreviations*: SG = Shotgun Sequencing, TE = Target enrichment
*References:* Bowtie2 (Langmead and Salzberg 2012); ngsLCA (Wang et al. 2022); CENTRIFUGE (Kim et al. 2016);

To date there is no executable aDNA workflow that relies on alignment-free methods for classification. The sedimix and aMeta pipelines use alignment-free classification to filter sequences before mapping or create a list of references for a subsequent alignment-based analysis. We found several studies that describe the use of alignment-free methods (see Tab. S1.1), but no general solution for the problem of false-positive taxa was described. E.g. Courtin et al. 2022 discuss the fact that they faced problems with false positive rates and recommend the use of alignment-based classifiers; Margaryan et al. 2018 state to “be cautious” with their taxonomic assignments and that Kraken should only be used as an exploratory step. In most cases, classification results are grouped on higher taxonomic levels (i.e. eukaryotes, Bacteria) or restricted to “expected” taxa (Courtin et al. 2022; Ottoni et al. 2019; Schulte et al. 2021).
Filtering strategies we selected for quicksand are further discussed in SI 4.

### Supplementary Section 2: Creating Simulated Datasets

This section describes the simulated datasets used for optimizing and benchmarking quicksand. An overview of the five datasets is provided in Table S2.1. Genomic sequences used for the simulations were downloaded from NCBI using NCBI Batch Entrez (https://www.ncbi.nlm.nih.gov/sites/batchentrez), the accession numbers are listed in Table S2.2 and Supplementary Table ST1.

**Table S2.1**
Overview of the five simulated datasets

| Dataset Size | Mammalian mtDNA | Mammalian nuclear DNA | Environmental DNA | Total number of reads |
|---|---|---|---|---|
| 0.1 | **500** | 24,500 | 475,000 | 500,000 |
| 1 | **5,000** | 20,000 | 475,000 | 500,000 |
| 10 | **50,000** | 25,000 | 425,000 | 500,000 |
| 100 | **500,000** | 0 | 0 | 500,000 |
| 1000 | **5,000,000** | 0 | 0 | 5,000,000 |

*Notes*: The columns show the number of simulated sequences in each dataset. The column "Dataset Size" refers to the number of simulated mtDNA sequences (in bold) relative to the number of mammalian mtDNA sequences in an average sedaDNA MiSeq dataset (see SI 2.1).

We simulated paired end (PE) Illumina MiSeq sequencing reads with universal Illumina sequencing adapters using gargammel `v1.1.2` with the flags `-ss MS` (Renaud et al. 2016). Gargammel is designed to simulate sequences from a single "endogenous" genome with the option to add bacterial contamination. To simulate metagenomic datasets, with sequences coming from multiple (mammalian) genomes, we treated all downloaded genomes as "bacterial contamination" by placing them as indexed fasta file in the `bact/` directory of the gargammel working directory and executing gargammel with the `--comp 1,0,0` flag.

Each of the five datasets was created three times independently, incorporating no, medium or high levels of aDNA deamination damage (C-to-T substitutions) in the terminal five base-pairs of each DNA sequence using the gargammel `-matfile` option. The matrix files for medium and high damage were the ones used for the verification of *vgan/euka* by Vogel et al. 2023 and are based on patterns observed in ~4000 years old neolithic samples (Günther et al. 2015; C-to-T substitution frequencies ranging from 1% at the 5th to 17% at the terminal base position) and ~7000 years old mesolithic samples (Olalde et al. 2014; 14-33%), respectively. The read length distribution, provided to gargammel with the `-f` flag followed the distribution observed in shotgun sequences of a Neanderthal sample from *Vindija Cave* in Croatia (Prüfer et al. 2017) with sequence lengths restricted to a minimum of 35 and a maximum of 100 base pairs (`--minsize 35` and `--maxsize 100` flags).

The creation of the simulated MiSeq PE sequencing reads was followed by Illumina adapter-removal and overlap-merging to full-length DNA molecules using leeHom (Renaud, Stenzel, and Kelso 2014) with the `--ancientdna` flag to allow partial overlap of simulated reads. The full-length sequences of the mammalian mtDNA, mammalian nuclear DNA and environmental background DNA were simulated for each dataset and damage profile independently and combined into FASTQ files using the unix 'cat' command.

#### 2.1 Mammalian mtDNA and Nuclear Genomes

Sequences were simulated from Mammalian families found in Pleistocene cave sediments, namely, Hominidae (Neanderthal), Elephantidae (Mammoth), Hyaenidae (Cave hyaena), Bovidae (Bos taurus) and Suidae (Sus scrofa) in proportions adopted from Slon et al. 2017 (see Table S2.2). The number of simulated mtDNA sequences increased across the five datasets from 500 to 5,000,000. The number of 5,000 mammalian mtDNA sequences was picked as baseline (Dataset Size 1) as it corresponds to the average number of mammalian mtDNA sequences in target-enriched sedaDNA datasets sequenced with ~500,000 reads (~4,800 sequences, for samples published in Slon et al. 2017).

**Table S2.2**
Mammalian genomes used as source for gargammel (NCBI accession numbers)

| Mammalian Family | mtDNA Genome | Nuclear Genome | Proportion |
|---|---|---|---|
| Hominidae | KC879692.1 | GCF_000001405.25 | 0.01 |
| Elephantidae | EU153448.1 | GCA_000001905.1 | 0.04 |
| Suidae | KC469586.1 | GCA_000003025.6 | 0.15 |
| Bovidae | DQ124371.1 | GCA_000003055.5 | 0.3 |
| Hyaenidae | JF894377.1 | GCA_008692635.1 | 0.5 |

*Notes*: The same proportion of sequences per genome was used in all dataset sizes.

#### 2.2 Environmental Background DNA

We simulated ancient DNA sequences from bacteria, archaea, fungi, plants and viruses, as proposed by Vogel et al. 2023 to explore the effect of ‘background DNA’ (DNA from sources that are not represented in the constructed reference database) on the quicksand results. Datasets with less than 500,000 mammalian mtDNA sequences were filled up with simulated background DNA as shown in Table S2.3. The full list of genomes used as source for the background DNA is provided in supplementary table T1.

**Table S2.3**
Number of “ background DNA” sequences simulated per dataset.

| Dataset Size | Bacteria | Archaea | Funghi | Plants | Viruses |
|---|---|---|---|---|---|
| 0.1 | 400000 | 35000 | 30000 | 8000 | 2000 |
| 1 | 400000 | 35000 | 30000 | 8000 | 2000 |
| 10 | 350000 | 35000 | 30000 | 8000 | 2000 |

Notes: See Supplementary Table ST1 for a list of reference genomes used

### Supplementary Section 3: Optimizing Database Kmer Size for sedaDNA Datasets

The optimal kmer length was determined using three criteria:

1. The *accuracy,* defined as the number of true positive sequence classifications divided by the total number of identified sequences (see main text)
2. *The sensitivity*, defined by the number of true positive sequence classifications (see main text)
3. The number of identified false positive families

#### 3.1 Number of False-positive families

We quantified the total number of false-positive family assignments in KrakenUniq (not quicksand as a whole) for each dataset, kmer size, and damage profile. Fig. S3.1 and S3.2 show that the number of false positive family assignments is primarily driven by the DNA damage profile (panels A, B and C) and the size of the simulated dataset. The kmer size only mattered when a unique kmer filter was applied to reduce taxonomic noise (see SI 4). In that case, smaller kmer-sizes show an increased number of false positives, because more unique kmers are generated from the same sequence, making it more likely to pass the filtering thresholds.

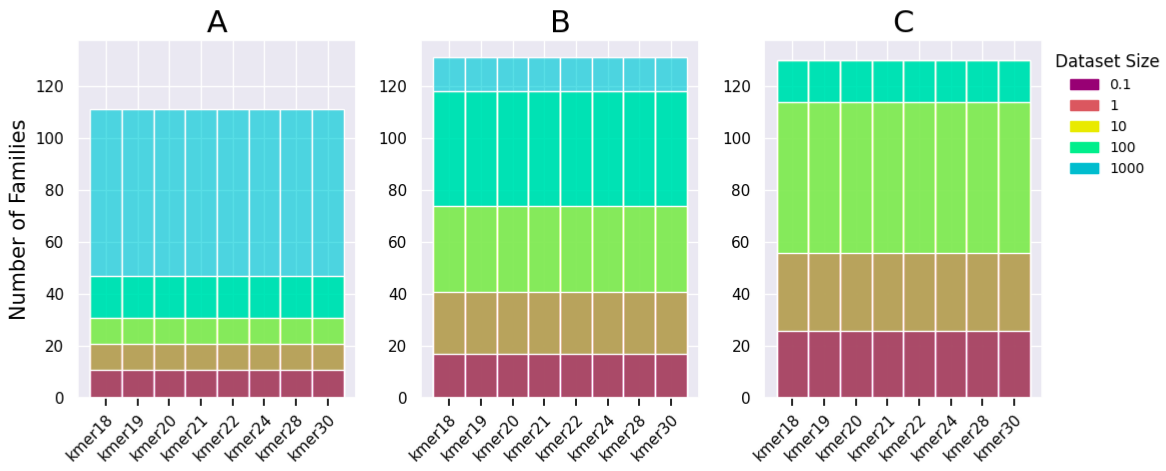


**Figure S3.1 |** Number of false positive **KrakenUniq** family assignments across different kmer sizes without the unique kmer-filter (see SI 4). Results for (A) undamaged, (B) medium damaged, and (C) highly damaged datasets. The barplots show the number of false-positive families found in each dataset with the respective kmer-size. Notably, 131 is the maximum number of mammalian families detectable in the database. Dataset size: see Tab. S2.1

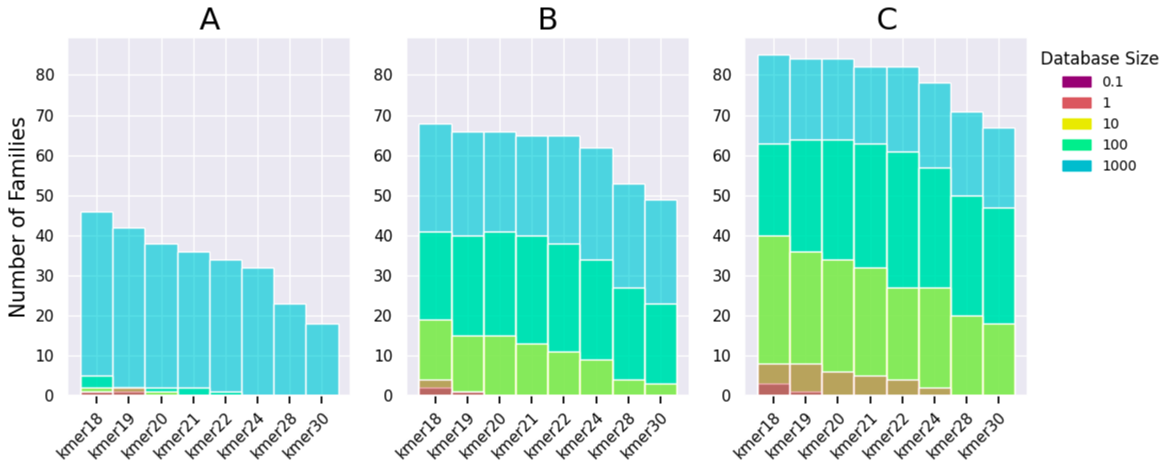


**Figure S3.2 |** Number of false positive **KrakenUniq** family assignments across different kmer sizes with an applied threshold of 129 unique kmers per family (see SI 4). See Fig. S3.1 for figure caption.

The high number of false-positive assignments is expected, as kmer-based methods are known to generate a substantial number of background identifications. It shows the importance of filtering KrakenUniq identifications and applying additional downstream filters to eliminate false-positive family assignments. We conclude that kmer size alone does not substantially influence the rate of false-positive assignments and that the counting of unique kmers is not sufficient as a standalone filter.

### Supplementary Section 4: Implementing Filters for the Removal of False-positive Family Assignments

The identification of false positive assignments is a well known problem for metagenomic analyses in general and for kmer-based classifiers in particular. To address this, several filtering strategies have been developed (see Table S1.1 for overview), which generally fall into the following categories:

1. **Quantity-based filters**: Filters that remove classifications which do not pass a predefined threshold (e.g minimum sequence counts or percentages of counts)
2. **Coverage-based filters:** Filters that assess the evenness of coverage in the alignment. Uneven coverage indicates mismapping, e.g. sequences clustering at conserved regions in the reference genome.
3. Incorporation of **biological expectations** - i.e. removal of *unlikely* species from the assignment based on known biological constraints.

We explored both quantity-based (SI 4.2) and coverage-based (SI 4.3) filters and established a set of recommended thresholds. However, most filters are not applied automatically within the pipeline. Instead, the relevant values for filtering are reported in the final summary report and need to be applied by the user. These statistics are calculated from the BWA alignments after PCR duplicate removal. We therefore aim to pass as many KrakenUniq assignments as possible to the mapping step.

However, KrakenUniq assigns sequences to all possible families in large and highly damaged datasets (Fig. S3.1). Forwarding all these assignments to the mapping step negates the speed advantage of kmer-based classification. In order to keep runtime low and to reduce the generation of non-informative (“junk”) output files, we apply a low unique kmer filter to remove false-positives prior to mapping (SI 4.1).

#### 4.1 Removal of KrakenUniq background identifications using unique kmers

In contrast to Kraken or Kraken2, KrakenUniq reports the number of unique kmers found for each taxonomic classification, a value that serves as a proxy for the breadth of coverage in a hypothetical alignment with the reference genome in the database. Filtering assignments by the number of unique kmers was proposed by the authors of KrakenUniq, who recommended a threshold of 1,000 kmers per species to eliminate “background identifications”, a threshold confirmed by the authors of aMeta (Pochon et al. 2022; and recently Oskolkov 2025). We believe that this threshold is too strict for quicksand - given that additional filtering steps are applied downstream, following actual sequence alignments.

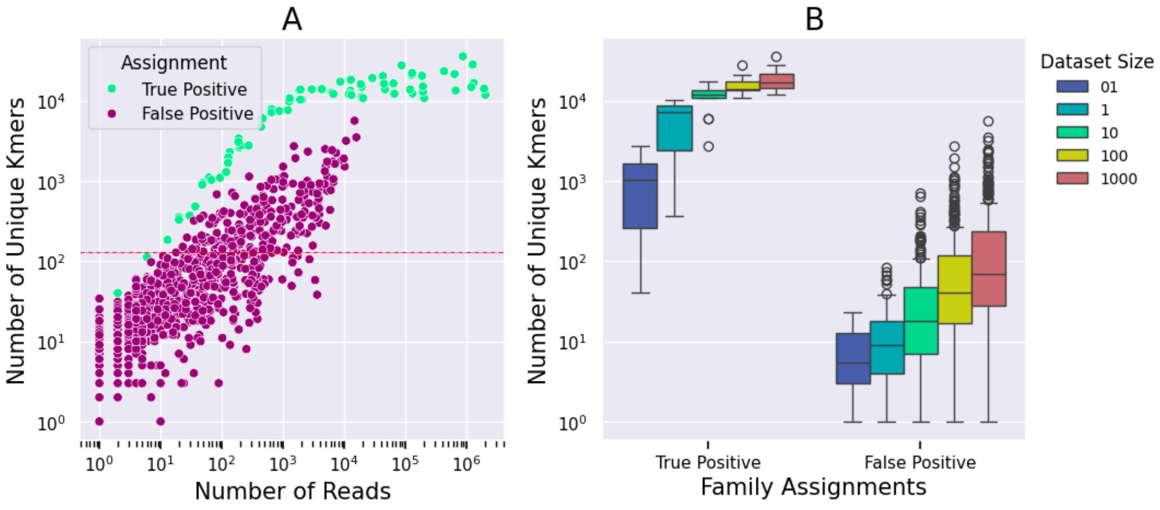


**Figure S4.1 |** Number of unique kmers assigned for each family in the simulated datasets (kmer-size 22). (A) By the number of reads, the red line indicates the default cutoff of 129 kmers. (B) By true and false positive family assignment, stratified by the dataset size (see Tab. S2.1). Boxplot formatting follows Fig.2;

In the simulated datasets, we observed that the number of unique kmers assigned to false-positive families increased with dataset size, rising from a median of 5 (max: 23) in the smallest to 69 (max: 5,616) in the largest dataset (Fig. S4.1B). This suggests that a percentage-based filtering threshold, e.g. one that removes families with fewer than 1% of unique kmers, is more effective as it does not depend on the size of the dataset. However, it should be noted that a percentage-based threshold may lead to the loss of true positive families especially in cases where the distribution of sequences across different families is highly uneven.

Families filtered from the KrakenUniq assignment *are not listed* in the summary report of quicksand (only in the KrakenUniq reports generated for each dataset). Therefore, to avoid the "silent" removal of potential true-positives, we implemented a minimally invasive static cutoff of 129 unique kmers, corresponding to ~150bp of covered sequences in the reference genome at a kmer-size of 22. This threshold successfully removed 100% of the false-positive families in the two smallest datasets and 75% and >50% in the second-largest and largest datasets, respectively. However, in the smallest dataset this filter also removed the Hominidae family (with five sequences), setting quicksand's default detection threshold to approximately ten sequences per biological family. While this threshold is fully customizable via the --krakenuniq_min_kmers flag (we discuss low-read sensitivity in SI 6), all subsequent filtering steps were tested with a unique kmer filter of 129 already applied.

#### 4.2 Minimum Percentage of unique Sequences per Family (PSF) Filter

Filters based on a minimum percentage of sequences assigned per biological family ("PSF") have shown to be effective in removing misidentified sequences (Tab S1.1). To explore the impact of this approach on the quicksand identifications, we compared the relative contribution of correctly and incorrectly identified families in the simulated data to the total number of mapped and deduplicated sequences (Fig. S4.2).

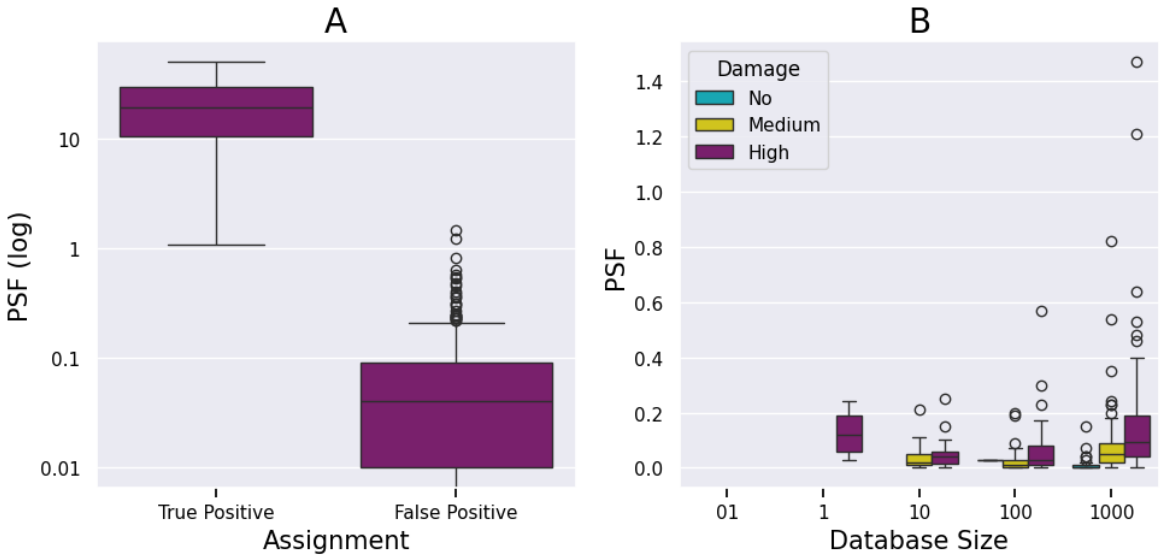


**Figure S4.2 |** Percentage of mapped and deduplicated sequences assigned to correctly and incorrectly identified taxa. (**A)** Distribution of PSF-values in true positive and false positive assignments (log-scaled). (**B**) Distribution of PSF-values in false positive taxa only, stratified by dataset size and damage profile. For dataset sizes, see Tab S2.1. Boxplot formatting follows Fig.2.

False-positive families are supported by a low PSF (median below 0.1%), but are generally more abundant in the larger and more damaged datasets. The PSF values are listed as 'FamPercentage' in the quicksand final summary report. We recommend a PSF threshold of at least **0.5%**.

#### 4.3 Minimum Proportion of Expected Breadth (PEB) Filter

As a second filter, we evaluated the evenness of coverage of mapped sequences along the reference genome for each family-level assignment. High evenness of coverage indicates that sequences are randomly distributed across the reference genome, which is expected when the source of the mapped DNA is closely related to the reference, such as from the same or a closely related species. In contrast, sequences that originate from more diverged sources may only map to conserved regions, resulting in clustered alignments and low evenness of coverage. Filtering based on evenness of coverage can therefore help identify and remove false positive taxonomic assignments.

The parameter reported in the quicksand final summary report for evaluating coverage evenness is the **proportion of expected breadth**. To understand this metric, it is necessary to define two key statistics calculated from an alignment:

- **Breadth of coverage**, defined as the proportion of the reference genome covered by at least one sequence.
- **Genomic coverage** (or depth of coverage), defined as the average number of times each base in the reference genome is covered by mapped sequences.

Under the assumption of random mapping to the correct reference genome, the breadth of coverage is a function of the genomic coverage and can be calculated using the formula empirically determined by Olm et al. 2021 (1).

$$(1)\ breadth\ of\ coverage\ =\ 1\ -\ e^{-0.883\ *\ coverage}$$

We refer to the calculated breadth of coverage as the **expected breadth of coverage**, as it assumes mapping to the correct reference genome. To evaluate deviations from this expectation, we calculated for each family the **proportion of expected breadth (PEB)**, defined as the ratio of the observed to the expected breadth of coverage. We found that the PEB was a suitable metric for distinguishing true from false positive family assignments (Fig. S4.3). For correct family assignments, the observed breadth of coverage matches the expectations (PEB around 1), while the false-positive families show PEB values between 1 and 0.2. We found that false-positives with low genomic coverage (<0.1×), match the expected breadth more often, presumably due to the small number of mapped sequences appearing randomly distributed, whereas only higher sequence counts reveal the clustering of sequences.

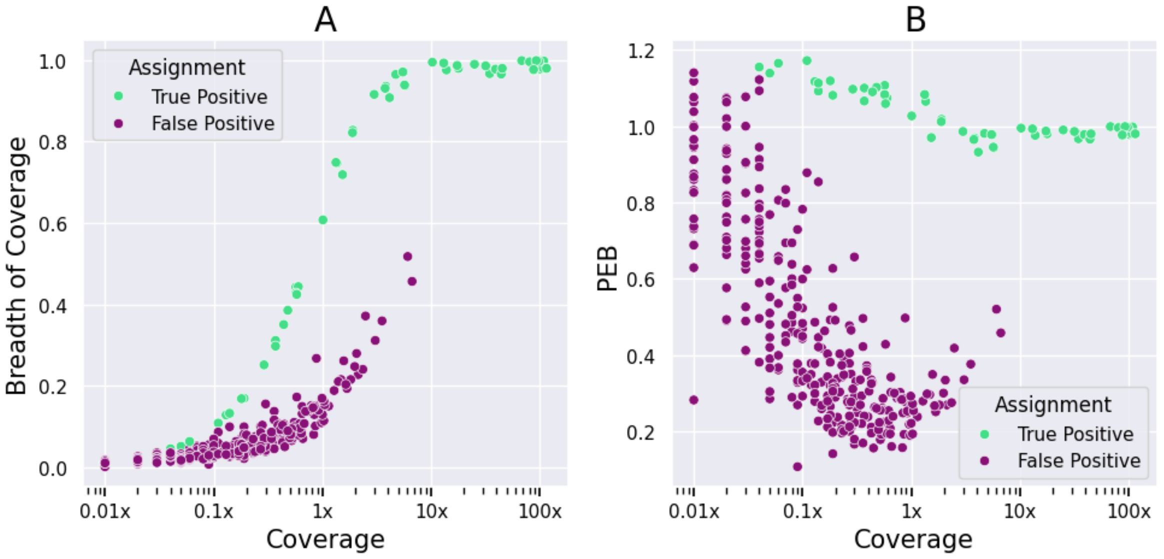


**Figure S4.3 |** True and False positive family assignments stratified by the genomic coverage of each alignment and (A) the breadth of coverage, (B) the proportion of expected breadth (PEB)

Based on the simulated datasets, we recommend removing family-level assignments with a proportion of expected breadth (PEB) below 0.5. Encouragingly, when applying the recommended filter cutoffs for both PSF and PEB, all false positive taxa were successfully removed from the quicksand results.

### Supplementary Section 5: Benchmarking quicksand against BLAST/MEGAN and euka-based Workflows

This section describes how the BLAST/MEGAN and the *vgan/euka*-based pipelines were executed for the benchmarking of quicksand.

#### 5.1 Running the BLAST/MEGAN-based Pipeline

We implemented a Nextflow version of the BLAST/MEGAN-based pipeline outlined in Slon et al. 2017 for the analysis of capture-enriched ancient mammalian and hominin mtDNA from sediments (https://github.com/merszym/blastmegan_nf). Briefly, sequences were assigned to a mammalian family using BLAST and MEGAN. For each detected family, sequences were re-mapped with BWA to all available reference genomes from that family. The alignment with the highest number of mapped sequences was then selected for downstream processing. As in quicksand, alignments were filtered (removing unmapped reads and those with mapping quality below 25) and PCR duplicates were collapsed into unique DNA molecules using bam-rmdup. Deamination frequencies were then calculated from the resulting alignments. The pipeline differs from the described workflow to allow the input of multiple already demultiplexed BAM files and to generate a summary report matching the one created by quicksand.

As the BLAST reference database, we used the same set of 796 complete mammalian mtDNA genomes described in Slon et al. 2017, which were downloaded from NCBI RefSeq in January 2016. These genomes were concatenated into a single FASTA file and indexed with BLAST. The ABIN file required for MEGAN (nucl_acc2tax-Nov2016.abin) was downloaded from the MEGAN6 website in 2016. For mapping with BWA, we compiled a set of mtDNA reference genomes (GENOMES) from NCBI RefSeq release 218 to ensure consistency with the references used for quicksand. We then executed the nextflow pipeline with the following command:

```
nextflow run merszym/blastmegan_nf --split INPUT --acc2taxid ABIN --database
FASTA --genomes GENOMES -profile singularity --pseudouniq_filterflag 1
--pseudouniq_mindup 1
```

We ran the BLAST/MEGAN pipeline with the flags --pseudouniq_filterflag 1 (remove only unmerged reads, keep unmapped reads) and --pseudouniq_mindup 1 (retain sequences that appear only once).

#### 5.2 Running the vgan/euka-based Pipeline

Direct performance comparisons between *vgan/euka* and quicksand were complicated by several factors. *vgan/euka* does not implement a PCR deduplication step, resulting in a higher number of total reads compared to quicksand. Additionally, *vgan/euka* assigns sequences to different taxonomic hierarchies. Hyena sequences

are assigned to the Carnivora pangenome subgraph (order level) but human sequences to the Homininae graph (at the sub-family level) (see Vogel et al. 2023, Supplementary Database Table for taxonomic group details). As a result, we focused on comparing only accuracy and runtime, rather than raw read counts, between *vgan/euka* and quicksand.
Since *vgan/euka* does not natively support batch processing of multiple files and requires input in FASTA format, we implemented an euka-based pipeline in Nextflow (https://github.com/merszym/euka_nf) for benchmarking quicksand. This pipeline added the BAM-to-FASTA conversion, enabled parallel processing of *vgan/euka* for multiple files, and compiled a final summary table for comparison with quicksand. We ran *vgan/euka* with default parameters, including the --outFrag flag, to get assignment statistics for each input sequence.

We downloaded the *vgan/euka* database (EUKADIR) as indicated in https://github.com/grenaud/vgan/wiki/euka#quick-start. The taxonomy (a folder containing the NCBI names.dmp and nodes.dmp files) required for the taxonomic level annotation (TAXONOMY) was downloaded in january 2025 from https://ftp.ncbi.nlm.nih.gov/pub/taxonomy/taxdmp.zip

For benchmarking, we ran the pipeline with the following command:

```
nextflow run merszym/euka_nf --split INPUT --euka_dir EUKADIR --taxonomy TAXONOMY -profile singularity
```

### Supplementary Section 6: Detecting the Hominidae family based on low sequence counts

As outlined in SI 4.3, we implemented a filter-threshold of 129 unique kmers for KrakenUniq family assignments to remove background identifications and to reduce runtime in the downstream mapping step. This threshold removed the true positive Hominidae assignment in the smallest simulated dataset with five human mtDNA sequences. Here we explored the effect of dropping the unique kmers filter for detecting families supported by low sequence counts.

Specifically, we ran additional simulations to address two questions:

1. How does quicksand perform when analyzing datasets consisting of fewer than 30 human sequences?
2. How reliable are low-read human assignments with another dominant mammalian family present in the sample?

#### 6.1 Analyzing Low-count Hominidae Sequences

We tested how consistently quicksand found true-positive families based on low sequence quantities. From the pool of 50,000 highly damaged simulated Neanderthal sequences (*Dataset size 1000, high damage profile; see SI 2*), we repeatedly (100 replicates each) subsampled between 5 and 30 random sequences into separate files (2600 in total) that were then analyzed using quicksand.

Without applying the unique kmers filter, quicksand was able to detect the Hominidae family in 90% of the subsamples with only five mtDNA sequences, and in 100% of subsamples with seven mtDNA sequences (Fig. S6.1C). However, not all false positive families were removed by the downstream filters. As shown before (see SI 4.3), families with low numbers of sequences escape PEB filtering, as the distribution of mapped reads looks sufficiently random to match the expectation of a true positive family. False positives (Cercopithecidae and Muridae), passing both the 0.5% PSF and the 0.5 PEB filter already appeared in replicates with as few as ten input sequences (Fig. S6.1A,B).

While only approximately 17% of the subsamples with 30 input sequences showed an ‘Ancientness’ rating of ‘+’ or ‘++’ (see main text), we observed stable mean terminal deamination rates (ranging from 25% to 35%, with 95% confidence intervals between 20% and 40%; Fig. S6.1C) across all subsamples with 16 or more sequences. These point estimates match the expected deamination rates of ~30% at both the 3′ and 5′ ends. Samples with fewer sequences exhibited greater variation in terminal deamination rates, with mean values ranging from 25% to 40% and 95% confidence intervals spanning from 15% to 50% for both 5′ and 3′ C-to-T deamination rates.

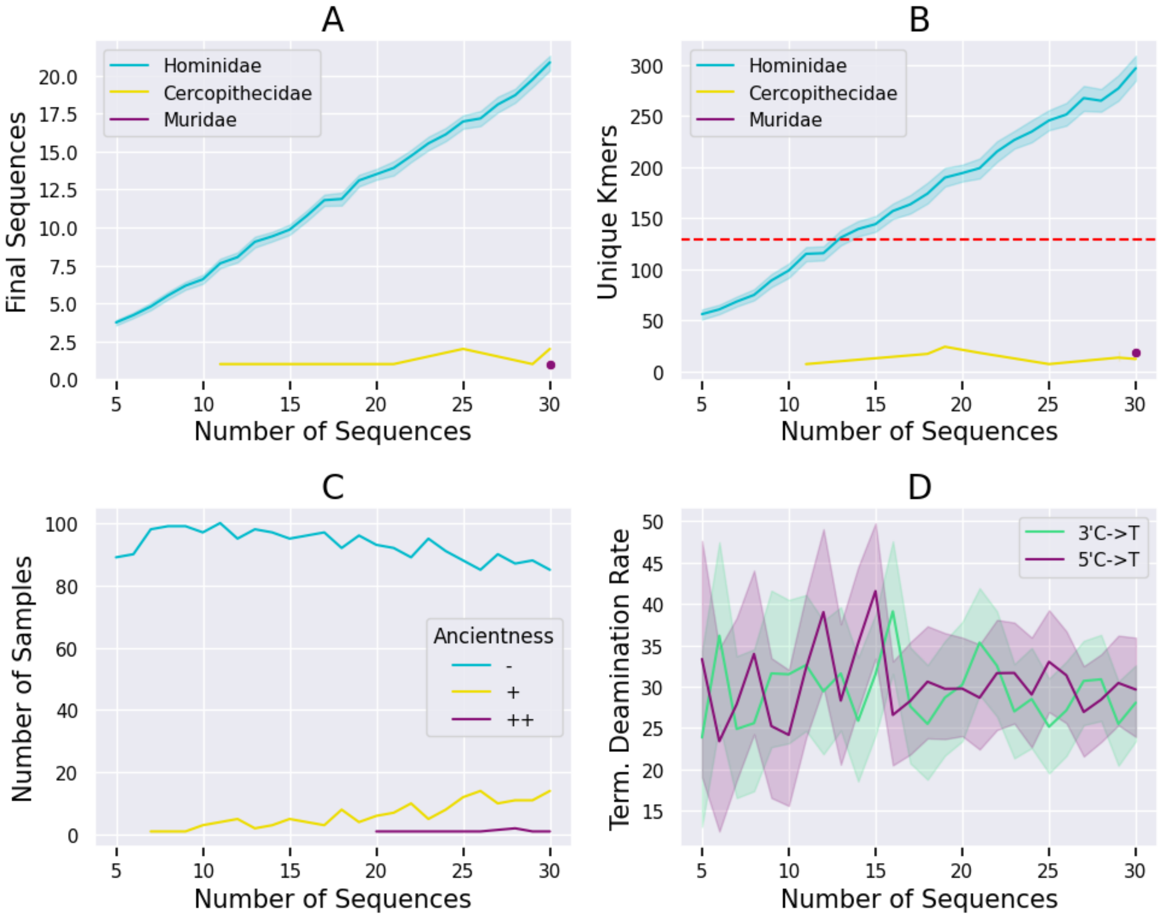


**Figure S6.1 | Results of family assignments with low sequence counts**. All plots are stratified by the number of Hominidae sequences put into quicksand (100 replicates each). (A) Mean number of final sequences (mapped, deduplicated, and bed-filtered) obtained for each assigned family. (B) Mean number of unique kmers obtained for each family. Red line indicates the default 129 unique kmer threshold. (C) Count of samples with the respective 'Ancientness' labels. (D) Mean 3′ and 5′ terminal C-to-T deamination rates (point estimates). Line plots in panel A,B and D show the mean (bold line) and the 95% confidence interval for repeated values (n = 100).

In summary, we show that quicksand is under some circumstances able to detect families even with as few as five input sequences. While the statistical power of these assignments is too low to label them as being 'Ancient' ('++' or '+'), detected deamination rates (point estimates) match the expected terminal deamination rates of the simulated data. We also note that the PSF filter and the PEB filter alone are not sufficient to remove false-positive assignments at these low read counts. However, lowering the unique kmer threshold from 129 to 50 could remove false positive families and improve detection in *low-yield samples* that would otherwise be classified as negative under the default settings.

#### 6.2 Analyzing Low-count Hominidae Sequences in the Presence of a High-count Hyaenidae Family

To test whether quicksand can detect low-count Hominidae sequences in the presence of a high-count Hyaenidae background, we added between 5 and 15 Hominidae sequences (randomly drawn as described in SI 6.1) to 2.5 million highly damaged Hyaenidae sequences previously simulated (*dataset size 1000; high damage profile;* SI 2), resulting in eleven datasets with identical Hyaenidae content but different numbers of random Hominidae sequences. We then analyzed these datasets using quicksand with default parameters, except with the unique kmer filter set to 0.

With the unique kmer filter omitted, all 131 mammalian families were identified by quicksand (Fig. S6.2 A). As previously reported (SI 4), these false-positive families represented between 0.01% and 5% of the "final" (mapped, deduplicated, and bed-filtered) sequences (Fig. S6.2C). Between 42 and 60 sequences were assigned to the Hominidae family, up to six times the number of human sequences actually present in the dataset (Fig. S6.2B).

quicksand groups the Hominidae family together with the false-positive family assignments, excluding them from downstream analyses when applying the default PSF and PEB filters (Fig. S6.2C). Notably, the Hominidae family would have passed the threshold for 129 unique kmer (Fig. S6.2D).

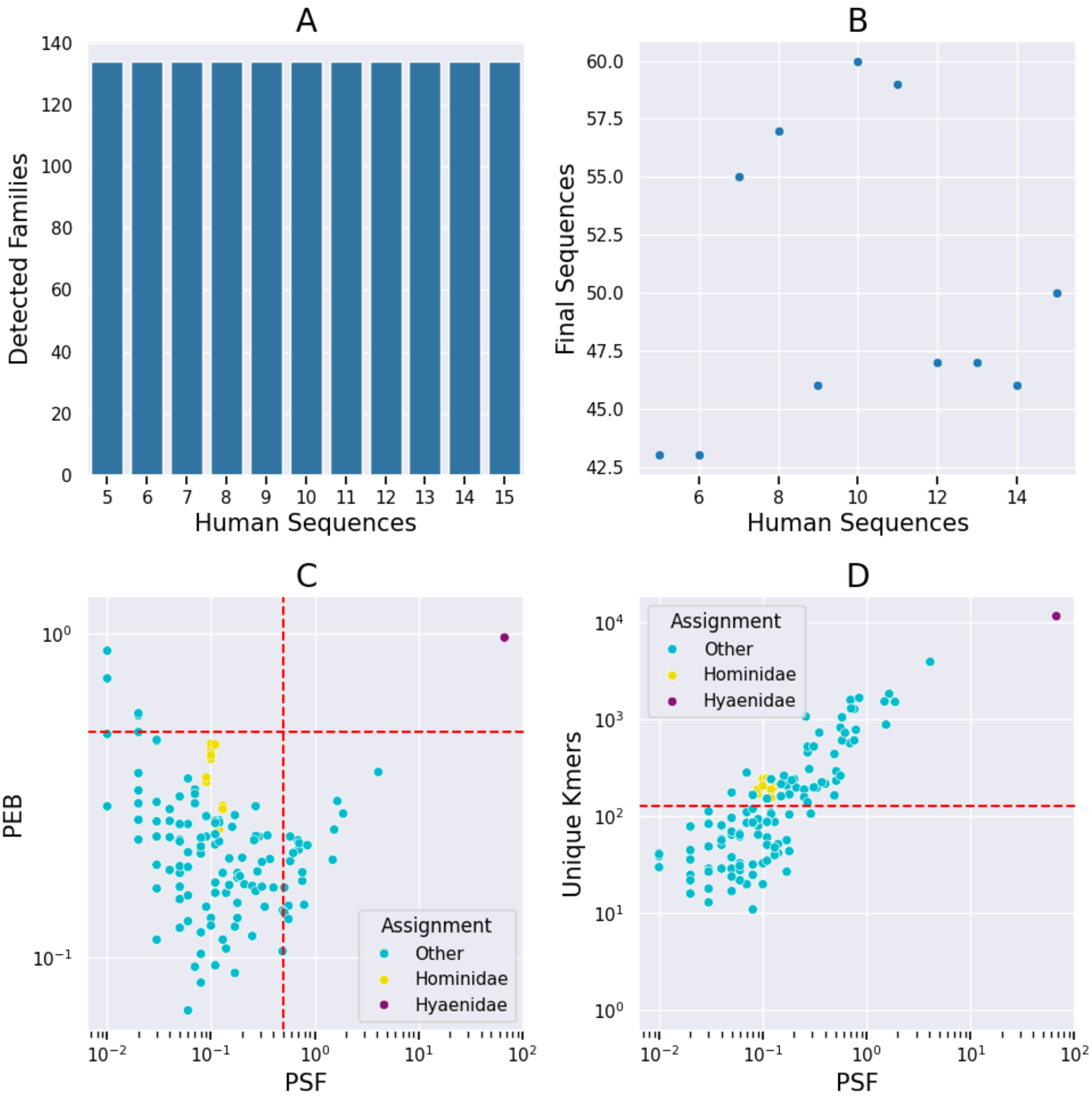


**Figure S6.2 | Hominidae assignments in the context of high-abundant Hyaenidae sequences**. Plots A and B are stratified by the number of Hominidae input sequences (one dataset per group), while plots C and D summarize all datasets. (A) Number of detected families in each dataset. (B) Final sequences (mapped, deduplicated, and bed-filtered) obtained for the Hominidae family assignment in each dataset. (C) Family assignments, stratified by the percentage of unique sequences per family (PSF) and their respective PEB. Red lines show the default quicksand filter thresholds for these two values. (D) Family assignments, stratified by PSF and the respective unique kmer count. The red line shows the default 129 unique kmer threshold.

We conclude that in the presence of an over-abundant mammalian family, the spill-over of sequences into the true-positive families prevents the detection of low-count true-positive families, as misclassified sequences dominate the respective assignments. As a result, such families are excluded as false-positive hits by the default quicksand filters.

While SI 6.1 suggests increased sensitivity when lowering the unique kmer filter, we consider a cutoff of 129 to be appropriate for the default quicksand execution, as it effectively removes a big portion of the KrakenUniq background identifications. If higher sensitivity is required, lowering the unique kmer cutoff (e.g., to 50 unique kmers) can be useful. However, this is only recommended for datasets in which mammalian sequences constitute only a small fraction of the total data (e.g., in shotgun-sequenced samples). In the presence of overly high-yielding mammalian families, low-count assignments should be handled with caution, as they are potentially a mix of true and false positive sequences and require further validation downstream.

### Supplementary Section 7: Analyzing sedaDNA from Denisova Cave, Main Chamber with quicksand v2.3

Work in Progress

We verified the performance of quicksand v2.3 and the kmer-size of 22 using a dataset published in Zavala et al. 2021 (see main text). In this section we show how the filter-thresholds we established with the simulated data perform on real data.

#### 7.1 Filtering the mammalian mtDNA capture libraries

We reanalyzed the 274 mammalian mtDNA capture libraries published as “initial screening” in Zavala et al. (2021). The mammalian capture panel used (AA75) was designed from 242 complete mammalian mtDNA genomes as a generic capture set intended to enrich samples for mammalian mtDNA fragments. This generic enrichment allows the assumption of a random distribution of mapped reads across the reference genome for each mammalian alignment, as required by the expectation-based PEB filter.

As determined for the simulated data, we applied both the 0.5% PSF filter and the PEB filter of 0.5 (Fig. S6.1). To assess the performance of these filters on samples from Denisova Cave, we assigned each detected family a label based on the likelihood of its DNA being present in the cave sediments of Denisova Cave: ‘Evidence’ if fossil evidence for the family exists at Denisova Cave (Vasiliev, Shunkov, and Kozlikin 2013; 2017), ‘Possible’ if the geographical distribution of the family overlaps with Denisova Cave, and ‘Unlikely’ if neither condition was met.

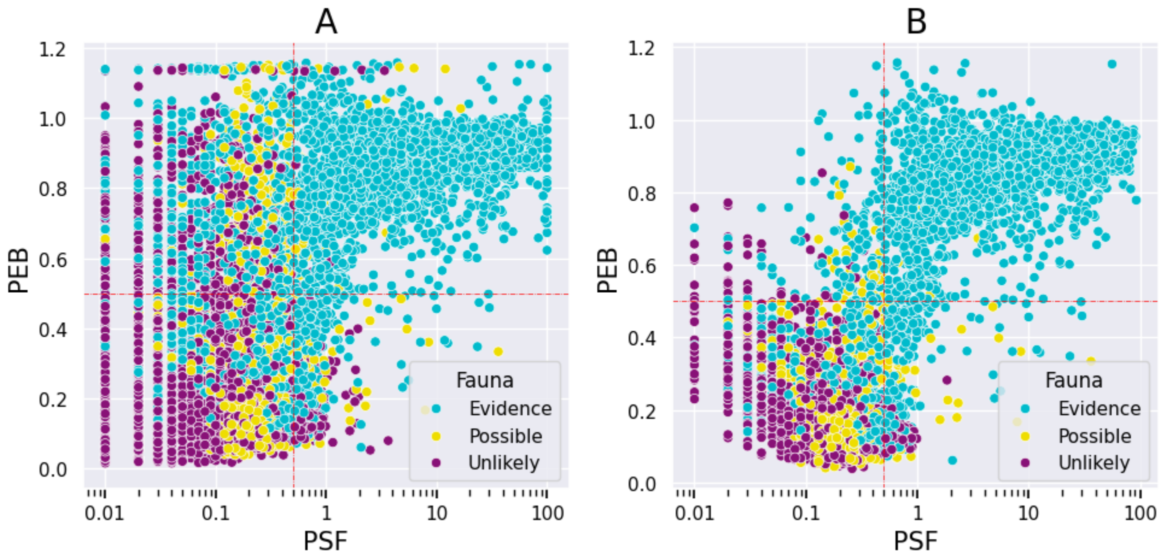


**Figure S7.1 | Effects of post-pipeline filters on quicksand results.** Each point in the scatterplots represents a mammalian family detected by quicksand v2.3 in one of the 274 mammalian mtDNA capture libraries, colored by the likelihood of its presence in Denisova

Cave. Families are stratified by the percentage of unique sequences in the sample (PSF) and the proportion of expected breadth (PEB) in the alignment to the respective “best” reference genome. Red lines indicate the applied filter thresholds; families falling below the horizontal line and to the left of the vertical line are removed from the results. (A) All family assignments. (B) Only families labeled as ancient (++) by quicksand.

The majority of “unlikely” or “possible” family classifications were assigned a PSF below 0.5% or a PEB below 0.5, matching the patterns of false positive assignments observed in the simulated datasets. As shown in Fig. S7.2, applying these filters results in only ten ancient families consistently meeting both thresholds, while additional seven families met both thresholds in some samples. Among the families meeting both thresholds, all (but one with a “Possible” label) carry the “Evidence” label, suggesting reliable assignments and successful filtering.

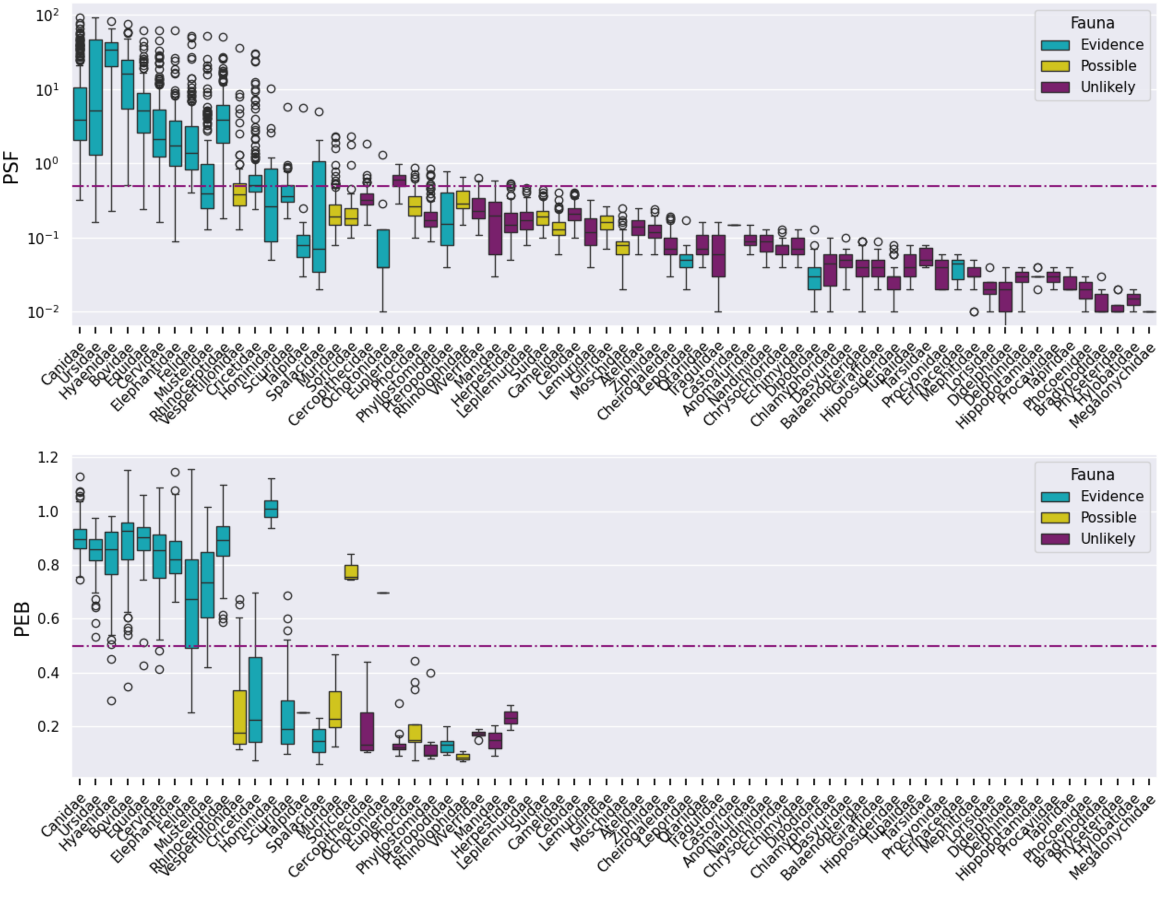


**Figure S7.2 | Filtering mammalian mtDNA enrichment samples.**
Both plots show the filtering steps for “ancient” families (++). **Upper panel:** Distribution of PSF values across all 274 samples and families. Box colors correspond to the likelihood classification of each family. The red horizontal line indicates the 0.5% PSF cutoff. **Lower plot:** Distribution of PEB values for the same family assignments that passed the 0.5% sequence cutoff. The red horizontal line indicates the 0.5 PEB cutoff. Boxplot formatting follows Fig.2.

After applying both filters, we found ancient mammalian mtDNA in 271 of the 274 sediment samples (Zavala et al. 2021: 167) and were able to detect additional ancient families in 205 samples that were not reported by Zavala et al. 2021. 44 samples showed fewer ancient families compared to Zavala et al. 2021 (Table S7.1).

**Table S7.1**
Number of samples with evidence for ancient DNA (++) of the listed mammalian families in both or either of the quicksand v2.3 and the Zavala et al. 2021 analyses.

| Family | both | only quicksand | only Zavala et al. 2021 |
|---|---|---|---|
| Canidae | 227 | 36 | |
| Ursidae | 207 | 43 | |
| Bovidae | 201 | 37 | 2 |
| Rhinocerotidae | 175 | 34 | 3 |
| Equidae | 169 | 14 | 3 |
| Cervidae | 136 | 56 | 3 |
| Hyaenidae | 134 | 58 | 1 |
| Felidae | 125 | 60 | 15 |
| Elephantidae | 112 | 59 | |
| Mustelidae | 45 | 23 | 3 |
| Hominidae | 4 | 6 | |
| Cricetidae | 1 | 16 | 13 |
| Vespertilionidae | 0 | 5 | 4 |
| Talpidae | 0 | 0 | 1 |
| Muridae | 0 | 0 | 1 |

#### 7.2 Lineage Assignment of the human mtDNA capture libraries

For all positive samples we performed a lineage assignment test as described in Meyer et al. 2016. The lineages for this test included Neanderthals, Hohlenstein-Stadel Neanderthals, the Sima de los Huesos hominins, Denisovans, the shared Denisova-Sima lineage, and modern humans. In brief, we counted for each sample the number of diagnostic positions in the rCRS genome overlapped by aligned mtDNA fragments and the number of positions where the fragments show the lineage-specific ('diagnostic') state for each lineage. The diagnostic positions for these lineages were taken from Zavala et al. 2021 and created by comparing multiple hominin genomes, selecting the sites where 99% of the genomes of one lineage differed from all others ("nochimp0.99", see Zavala et al. 2021: Supplementary Data File 3 for the used genomes). The support for each lineage was calculated as the percent of shared states among all overlapping diagnostic positions. Lineage assignments were retained if the lineage support exceeded 10% and at least three different shared diagnostic positions were identified. Diagnostic positions that were overrepresented in the dataset (i.e., present in more samples than two standard deviations above the mean number of samples per diagnostic position, see Zavala et al. 2021; SI 4.1) were removed and assignments to the modern human lineage were retained if they could be confirmed using only deaminated mtDNA fragments to reduce the impact of modern human contamination.

Vasiliev, Sergei, Michael Shunkov, and Maxim Kozlikin. 2017. "Megafaunal Remains from the Eastern Chamber of Denisova Cave and Problems of Reconstructing the Pleistocene Environments in the Northwestern Altai // Остатки Мегафауны Из Восточной Галереи Денисовой Пещеры И Вопросы Реконструкции Природной Среды Северо-западного Алтая В Плейстоцене". *Problems of Archaeology,*

*Ethnography, Anthropology of Siberia and Neighboring Territories - Novosibirsk* XXIII: 60–64.

Vogel, Nicola Alexandra, Joshua Daniel Rubin, Mikkel Swartz, Juliette Vlieghe, Peter Wad Sackett, Anders Gorm Pedersen, Mikkel Winther Pedersen, and Gabriel Renaud. 2023. “Euka: Robust Tetrapodic and Arthropodic Taxa Detection from Modern and Ancient Environmental DNA Using Pangenomic Reference Graphs”. *Methods in Ecology and Evolution* 14 (11): 2717–27. https://doi.org/10.1111/2041-210x.14214.

Wang, Yucheng, Mikkel Winther Pedersen, Inger Greve Alsos, Bianca De Sanctis, Fernando Racimo, Ana Prohaska, Eric Coissac, et al. 2021. “Late Quaternary Dynamics of Arctic Biota from Ancient Environmental Genomics”. *Nature* 600 (7887): 86–92. https://doi.org/10.1038/s41586-021-04016-x.

Wang, Yucheng, Thorfinn Sand Korneliussen, Luke E. Holman, Andrea Manica, and Mikkel Winther Pedersen. 2022. “ngsLCA—a Toolkit for Fast and Flexible Lowest Common Ancestor Inference and Taxonomic Profiling of Metagenomic Data”. *Methods in Ecology and Evolution* 13 (12): 2699–2708. https://doi.org/10.1111/2041-210x.14006.

Willerslev, Eske, Anders J. Hansen, Jonas Binladen, Tina B. Brand, M. Thomas P. Gilbert, Beth Shapiro, Michael Bunce, Carsten Wiuf, David A. Gilichinsky, and Alan Cooper. 2003. “Diverse Plant and Animal Genetic Records from Holocene and Pleistocene Sediments”. *Science* 300 (5620): 791–95. https://doi.org/10.1126/science.1084114.

Xu, Jierui, Elena I. Zavala, and Priya Moorjani. 2025. “Sedimix: A Workflow for the Analysis of Hominin Nuclear DNA Sequences from Sediments”, March. https://doi.org/10.1101/2025.02.28.640818.
